# Functional Implications of the Conformational Landscape of a Multidrug Transporter Revealed by Structures of Zebrafish Abcb4

**DOI:** 10.1101/2025.05.30.656806

**Authors:** Jingyu Zhan, Chao-Ming Hsieh, Lothar Esser, Zabrina C. Lang, Abraham J Morton, Robert Robey, Fei Zhou, Suresh V. Ambudkar, Rick K. Huang, Michael M. Gottesman, Di Xia

## Abstract

The hallmark of multidrug resistance conferred by human ABC transporter ABCB1 (hP-gp) is the recognition and efflux of diverse range of drugs, though the precise mechanism of polyspecificity remains unresolved. In aquatic animals such as zebrafish, Abcb4, a functional homolog to hP-gp, plays a vital role in surviving environmental toxicants. Here, we show that Abcb4 exhibits comparable basal and drug-stimulated ATPase activity to hP-gp. Using cryo-EM, we captured five inward-facing Abcb4 conformations with varying separations between its two lobes, illustrating its open-and-close motion. The range of separation exceeds that seen in published P-gp structures that appear to be conformationally restricted. This global open-and-close motion is coupled with individual helix movement, resulting in a highly fluid substrate-binding pocket. These dynamic changes, likely underlying the polyspecificity of substrate recognition, predict unconventional protein-ligand interactions that are supported by structures of Abcb4 bound to the P-gp inhibitors tariquidar and elacridar, and the substrate vincristine.

## Introduction

ATP-binding cassette (ABC) transporters, one of the largest membrane protein superfamilies, are found across all kingdoms of life and facilitate the transport of endogenous and exogenous compounds across cellular membranes^1^. ABC transporters share a common modular architecture consisting of two lobes; each is made of a conserved intracellular nucleotide-binding domain (NBDs) and a transmembrane domain (TMD) (Fig. 1A). ATP hydrolysis by the NBDs drives conformational changes of the TMDs, enabling substrate binding and release on opposite sides of the membrane^2^. In humans, ABC transporter malfunctions can lead to complex diseases, including cystic fibrosis^3^ and several liver and retinal disorders^4–7^. Additionally, overexpression of these transporters, especially by cancer cells, is linked to the development of multidrug resistance in cancer (MDR)^5,8^.

**Figure 1.**
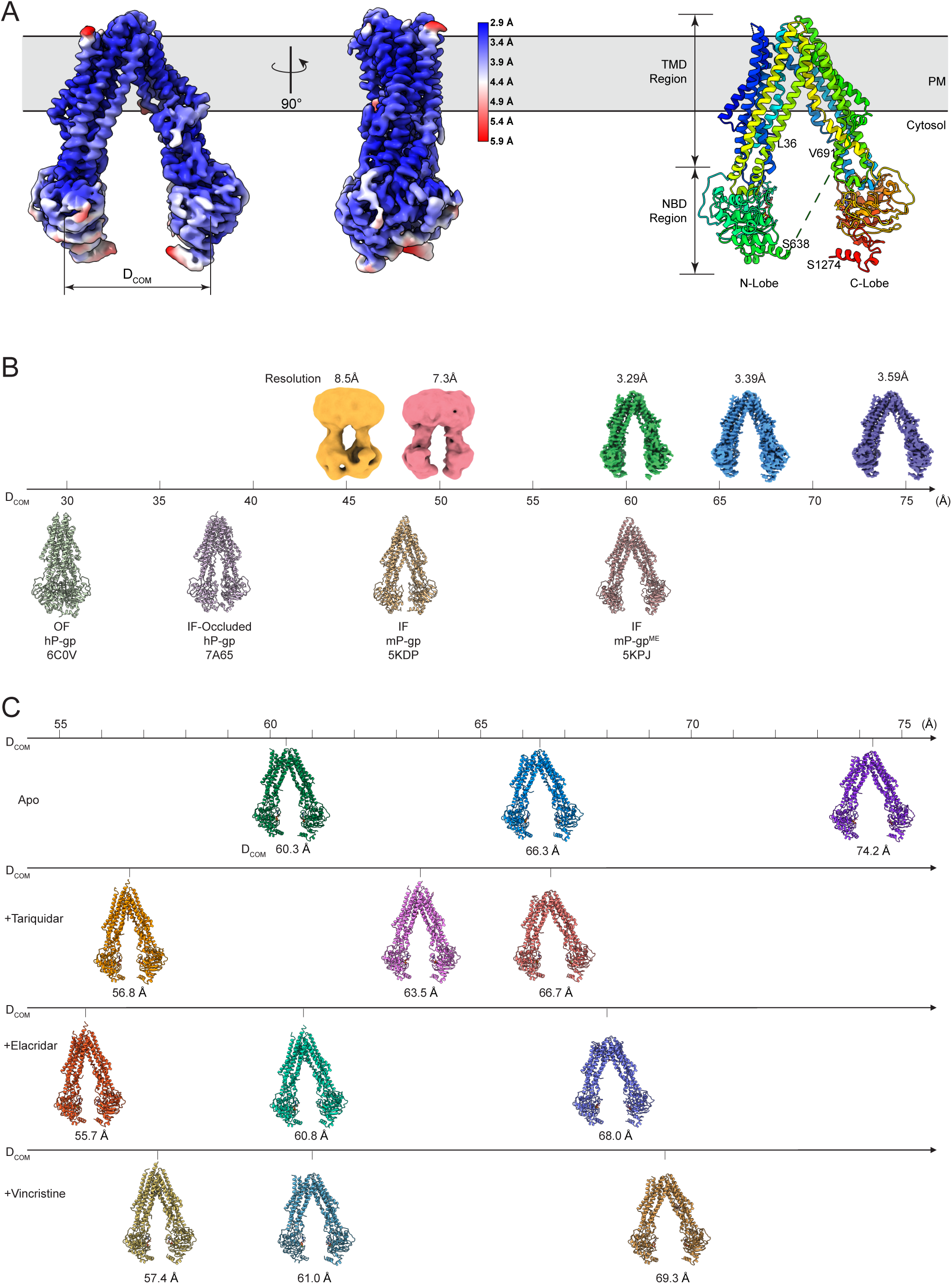
Cryo-EM structures and conformations of DrAbcb4 and its complex with drugs. (A) EM density in two orthogonal orientations and its derived ribbon model of DrAbcb4 in the IF^Nar^ conformation. Local resolution of the EM density is color-coded as shown in the vertical strip. The two horizontal lines delineate the TM region of the transporter. (B) EM density maps for the five classes of DrAbcb4 dataset are arranged by their D_COM_ values. Map resolutions are indicated. As references, the structures of hP-gp (PDB:6C0V) in OF conformation, hP-gp (PDB:7A65) in the IF occluded conformation embedded in nanodisc and bound with MRK16, the mP-gp (PDB:5KDP) in IF conformation confined in crystal lattice, and the methylated m-Pgp (PDB:5KPJ) in IF conformation embedded in crystal lattice are shown with their respective D_COM_ values indicated. (C) Ribbon models of Abcb4, built into three high-resolution reconstructions for each dataset, are shown as a function of D_COM._ From top to bottom: drug-free DrAbcb4, Abcb4 with tariquidar-, elacridar-, and vincristine-bound.

MDR conferred by ABC transporters is achieved through active efflux of a wide range of drugs from cells, reducing intracellular drug concentration and lowering drug efficacy. ABCB1, also known as MDR1 or P-glycoprotein (P-gp), is the first ABC transporter identified for its role in conferring MDR in mammalian cells including cancer cells^9^ and remains the most extensively studied. Beyond its role in MDR, P-gp exists in physiological barriers like the blood-brain barrier (BBB), blood-placenta barrier (BPB), and intestinal epithelium, protecting vital organs from xenobiotics while limiting therapeutic drug penetration^10^. P-gp and other ABC multidrug transporters, such as ABCG2 and ABCC1, also play key roles in the excretion of drugs from the liver into the bile, and from the kidney into the urine^8^. This protective role is evident in P-gp knockout mice, which, though phenotypically normal and fertile, exhibited hypersensitivity to chemicals and deficiency in the BBB^11,12^. A key objective in P-gp research is to understand the mechanism underlying its substrate polyspecificity, which may be explored effectively through structural studies. Indeed, numerous structures obtained via X-ray crystallography and cryo-EM have revealed remarkable versatility of P-gp in handling substrates (see reviews^2,13,14^). However, most drug-bound structures of P-gp were obtained under a small number of specific conditions, resulting in their clustering into a few discrete inward-facing (IF) conformations. These conformations are constrained by factors such as binding to specific monoclonal antibodies, incorporation into nanodiscs, crystal lattice packing, or covalent attachment to substrates^15–21^. They represent only a subset of conformational states that P-gp can adopt to handle a wide range of substrates.

Modeling P-gp function *in vivo*, for example in the blood-brain barrier (BBB), has shifted recently to organisms such as zebrafish, which possess a variety of ABC multidrug transporters homologous to mammalian transporters^22,23^. Compared to human, aquatic animals, constantly exposed to waterborne chemicals and toxicants, require conceivably more efficient detoxification mechanisms throughout their life cycle. Indeed, zebrafish have a genome that encodes a higher number of ABC efflux transporters including a P- gp ortholog, named Abcb4, which is broadly distributed in its embryo, adult BBB, and digestive system similar to human P-gp^24–26^. Unlike human ABCB4, which primarily functions as a lipid translocator, zebrafish Abcb4 (DrAbcb4) shares functional homology with human P-gp (hP-gp), including a virtually identical substrate specificity profile and 64% sequence identity^11,24,26^. Thus, zebrafish serves as a promising vertebrate model for studying P-gp function, including drug transport across the BBB and MDR *in vivo*.

Using wild-type DrAbcb4, we demonstrate its utility as a model system for structural and mechanistic studies of P-gp. Our analysis reveals multiple distinct IF conformations with very different separation distances between the two lobes of DrAbcb4, suggesting it can reach an extended range of conformations necessary for function. These conformations likely represent snapshots of DrAbcb4 undergoing spontaneous open-and-close motions, an essential process for the transport function of P-gp^27,28^. Moreover, this open-and-close motion is correlated with a complex movement of individual transmembrane helices (TMHs), which involves rotation, tilting, twists, unwinding, and even translational movement. This dynamic coupling alters the surface topography of the substrate-binding pocket encompassed by TMHs within the cell membrane. A correlative analysis reveals clustering of published P-gp structures, which is indicative of their conformations confined by crosslink modifications, by crystal contacts, by conformation-selective antibodies, or by nanodisc embedment. By contrast, Abcb4 exhibits a broader range of conformations that are accessible to substrates as shown by structures of Abcb4 bound to tariquidar, elacridar, and vincristine. Notably, we found that binding of these compounds, rather than nucleotide, promotes the two lobes moving closer. The fluid substrate-binding environment permits not only binding of different substrates but also the same substrate in different poses and may underpin the polyspecificity that allows P-gp to recognize and transport a wide range of substances.

## Results

### Zebrafish Abcb4 exhibits a similar ATPase activity profile as P-gp

We expressed recombinant DrAbcb4 in *Pichia pastoris* and purified it to homogeneity. The protein displayed monodispersity in solution under the purification conditions, as confirmed by size-exclusion chromatography (Supplementary Figure 1A) and electron microscopy (Supplementary Figure 2A). Additionally, the purified protein cross-reacted with the monoclonal antibody C219, known to recognize various mammalian P-gps^29^ (Supplementary Figure 1A, inset), which is consistent with previous reports^24,26^.

To further assess the activity of DrAbcb4, we measured its basal and drug-stimulated ATPase activity in both detergent micelle and lipid-detergent bicelle environments (Supplementary Figure 1B). The basal ATPase activity of DrAbcb4 was approximately 10.6 nmol P_i_/min/mg protein in dodecyl-D-maltoside (DDM) micelles and 5.9 nmol P_i_/min/mg in lipid bicelles, comparable to those of human and murine P-gp as previously reported^26,27^. Verapamil modulates the ATPase activity in a concentration-dependent manner, displaying a characteristic skewed bell-shaped curve. At lower verapamil concentrations, ATPase activity is stimulated up to 2-fold in the detergent solution and approximately 15-fold under the lipid bicellar conditions, whereas at higher concentrations it changes to inhibition (Supplementary Figure 1B). These results confirm that DrAbcb4 exhibits similar basal and verapamil-modulated ATPase activity to human and murine P-gp.

A comparable analysis with vincristine, a Vinca alkaloid antimicrotubule agent used in cancer therapy, showed little ATPase stimulation (Supplementary Figure 1C), which is consistent with a previous report using recombinant DrAbcb4. Tariquidar and elacridar, both third-generation inhibitors of P-gp known to suppress its ATPase activity^19^, have also been reported to effectively block the transport activity of DrAbcb4 in human HEK293 cells^26^. When the isolated, *Pichia*-expressed DrAbcb4 was tested for ATPase activity under detergent conditions, elacridar showed moderate inhibitory effect while tariquidar did not have much impact (Supplementary Figure 1C). These findings are consistent with previous reports showing that DrAbcb4 shares a substrate/inhibitor specificity profile with hP-gp^24–26^, further supporting its suitability as an *in vitro* model for structural and mechanistic studies of P-gp.

### Conformational landscape of wildtype zebrafish Abcb4

Characterization of purified DrAbcb4 demonstrated that the recombinant protein is monodisperse in solution, biochemically active, and suitable for structural studies. We aimed to obtain the structure of DrAbcb4 without constraints imposed by mutations, nanodisc embedding, antibody trapping, or containment in a crystal lattice. Using cryo-EM, we analyzed DrAbcb4 in the presence of ATPγS and Mg^2+^. This yielded five inward-facing (IF) conformations (Fig. 1B, Supplementary Figure 2, and Supplementary Table 1), each with a distinct separation distance (D_COM_) defined as the distance between the center of mass of the two NBDs^27^. These conformations likely capture snapshots of DrAbcb4 molecules undergoing the open-and-close motions in solution.

The three predominant classes were reconstructed to high resolution (3.29Å, 3.39Å, and 3.59Å, respectively, Fig. 1B and Supplementary Table 1), revealing D_COM_ values of 60.6Å (IF^Nar^), 66.3Å (IF^Med^), and 74.2Å (IF^Wide^), respectively. Notably, these D_COM_ values are larger than those for the structures previously reported (Table 1). The two minor classes with narrower D_COM_ separations did not reach high resolution because of lower particle populations representing more transient states. In the three higher resolution structures, the TM regions were consistently better resolved than the NBDs, displaying higher local resolutions and allowing for *de novo* model building (Fig. 1A, Supplementary Figures S2B, S3A and S3B). The NBDs were modelled by docking the homologous NBD models from murine P-gp (mP-gp, PDB:5KO2) followed by sequence substitutions and manual refinement based on EM density (Supplementary Figure 3C). Although the overall quality of EM density for NBD1 and NBD2 were lower, the density around the nucleotide-binding sites were well defined, allowing confident placement of bound ATPγS/Mg^2+^ (Supplementary Figure 3D).

**Table 1A.**
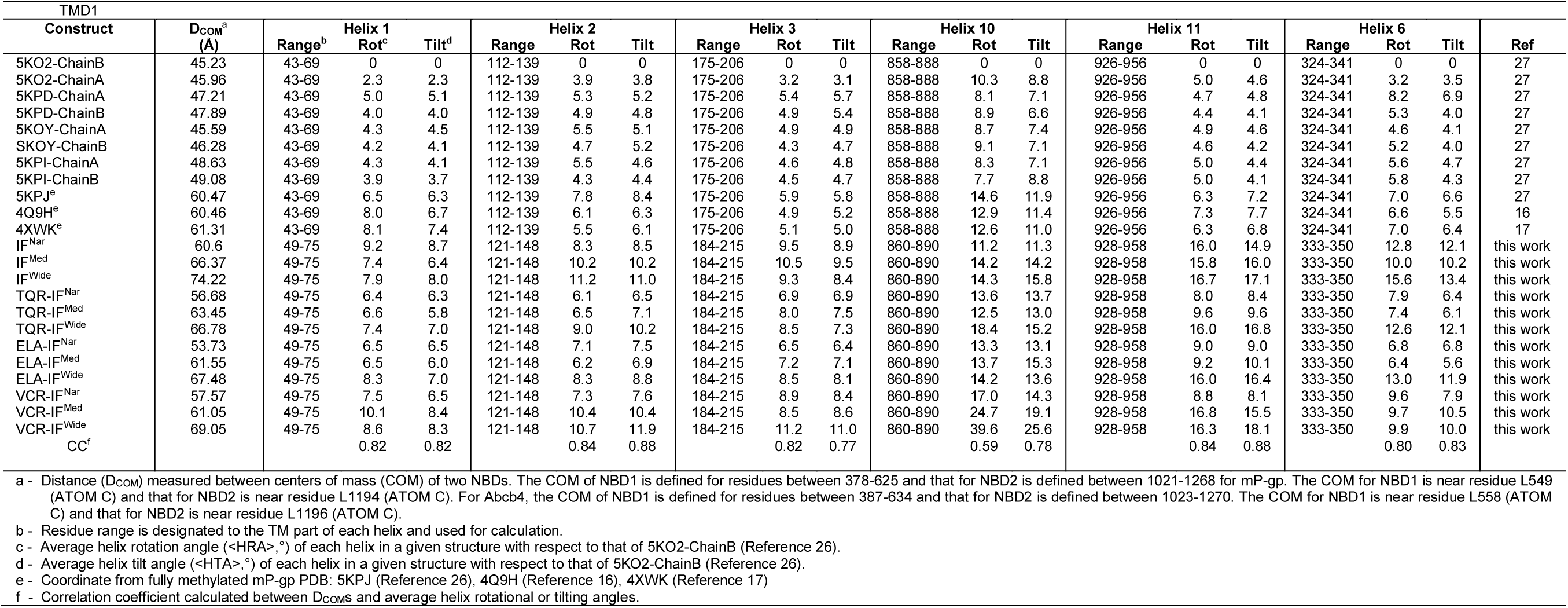
Correlation of the separation distances (DCOM) between the two NBDs with the magnitude of movement of individual helices for TMD1.

**Table 1B.**
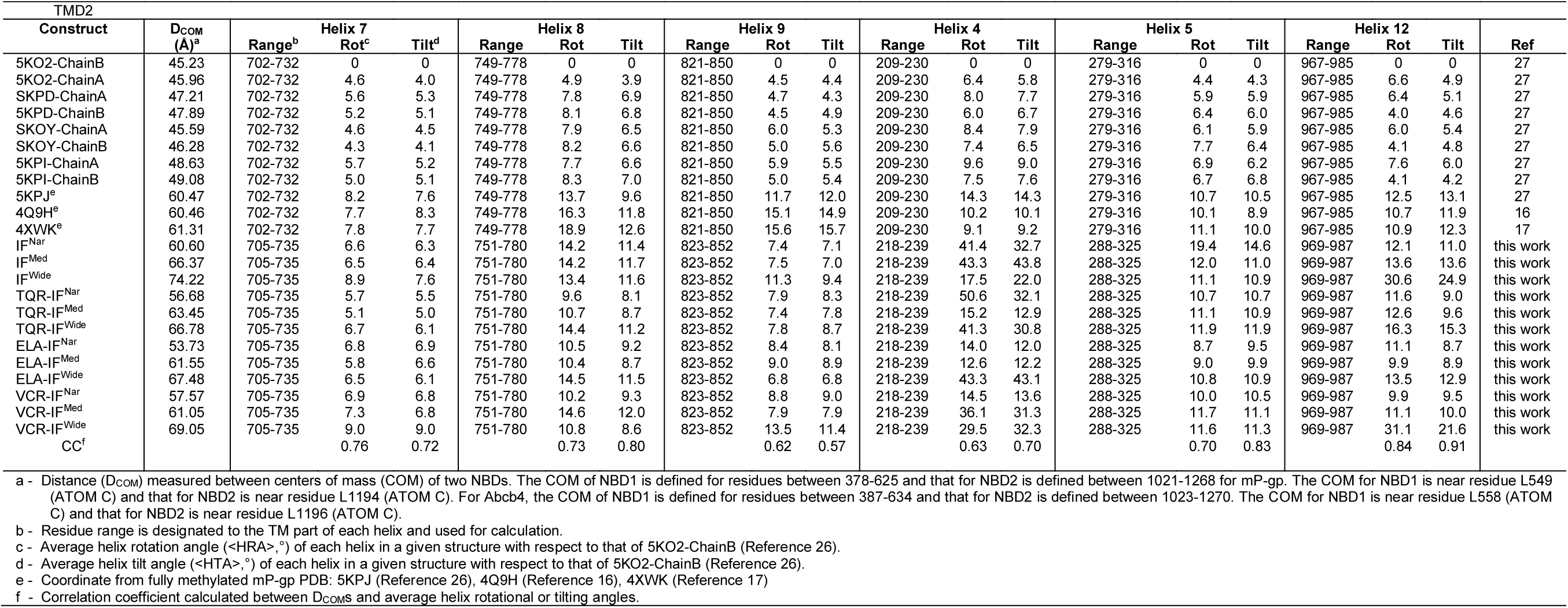
Correlation of the separation distances (DCOM) between the two NBDs with the magnitude of movement of individual helices for TMD2.

The overall architecture of the zebrafish Abcb4 closely resembles that of P-gp, featuring a core structure with two distinct lobes (Fig. 1A). Each lobe comprises a transmembrane domain (TMD1 or TMD2) and a nucleotide-binding domain (NBD1 or NBD2). Here, the domains are defined based on structure instead of sequence. Like human and murine P-gps, the linker (F636-K690) connecting NBD1 to TMD2 remains disordered, even though it is seven residues shorter than that of mP-gp or hP-gp (Supplementary Figure 4). Each TMD consists of six transmembrane helices (TMHs), with four helices from one half of its sequence and two domain-swapped helices (TMHs 4 and 5 in TMD2 and TMHs 10 and 11 in TMD1) from the other half. In all the three major states, both NBDs contain densities that can be confidently modeled with ATPγS/Mg^2+^ bound and remain well separated. This suggests that nucleotide binding alone is incapable of inducing NBD-dimerization in wild-type DrAbcb4 in the absence of substrate, at least not to a degree detectable by cryo-EM under our conditions. By contrast, structures of mP-gp with ATP/Mg^2+^ (PDB:8AVY, L335C+E/Q mutations) in detergent solution^21^ and hP-gp with ATPγS/Mg^2+^ (PDB:9CTG) embedded in nanodisc^30^, both in outward facing (OF) conformation, were obtained in the absence of substrate.

A nearly complete atomic model of DrAbcb4 was constructed, except for the two regions: the extracellular loop 1 (ECL1, residues 80-114), which connects TMH1 and TMH2, and the linker (residues 639-690) connecting the two lobes (Fig. 1). Importantly, the absence of the linker density in DrAbcb4, as well as in other full-length or linker-shortened P-gp structures, is likely due to their inherent flexibility^27^. Despite being unresolved structurally, the linker is essential for transport function. Any alterations that rigidify or shorten this region in P-gp result in inactive transporters, highlighting that the ability of the two lobes to open wide is required for its biological function.^27,28^

### Structures of DrAbcb4 in the presence of drugs

We aimed to examine structural changes in DrAbcb4 upon binding to specific drugs under solution conditions identical to those in the drug-free state. We selected vincristine (VCR), tariquidar (TQR), and elacridar (ELA), as DrAbcb4 confers resistance to vincristine in zebrafish embryos through its efflux activity^24^ that can be reversed by tariquidar or elacridar^26^. Moreover, the use of these compounds enables direct comparison to hP-gp, because structures of hP-gp bound to these drugs, as well as in apo form, have been reported in complex with the Fab fragment of a monoclonal antibody MRK16^19^. Notably, these hP-gp structures exhibited an IF conformation with a D_COM_ of approximately 38 Å, in which the substrate-binding pocket, whether empty or drug bound, is enclosed from the cytosolic side by kinked TMH4 and TMH10, defined as the IF occluded state^19^ (see PDB:7A65 in Figure 1B).

Following a similar workflow as in the study of the drug-free DrAbcb4, we acquired cryo-EM data sets of DrAbcb4 in saturation with tariquidar, elacridar, and vincristine (Supplementary Figures 5, 6, and 7). Interestingly, reconstructions of the DrAbcb4/TQR and DrAbcb4/ELA datasets reached higher resolutions than the drug-free DrAbcb4, while those from the DrAbcb4/VCR dataset had relatively lower resolutions (Supplementary Table 1), which suggests a potential stabilizing role of inhibitors, but not substrates, to the conformations of DrAbcb4 in solution. Three major conformations with different D_COM_ separations were obtained for each compound: DrAbcb4/TQR at 56.8Å, 63.5Å, and 66.7Å (designated as TQR-IF^Nar^, TQR-IF^Med^, and TQR-IF^Wide^, respectively); DrAbcb4/ELA at 55.7Å, 60.8Å, and 68.0Å (designated as ELA-IF^Nar^, ELA-IF^Med^, and ELA-IF^Wide^, respectively); DrAbcb4/VCR at 57.4Å, 61.0Å, and 69.3Å (designated as VCR-IF^Nar^, VCR-IF^Med^, and VCR-IF^Wide^, respectively) (Supplementary Table 1 and Table 1). The overall separations (D_COM_) observed in the presence of inhibitor and substrate were consistently narrower compared to those of the apo dataset (Table 1, Figure 1C), indicating that drug-binding may induce closing of the two lobes.

### Coordinated structural changes of DrAbcb4 with its open-and-close motions

The twelve high resolution structures of DrAbcb4, obtained either in the absence or presence of drugs, showed various separation distances (D_COM_) of the two lobes, representing snapshots of DrAbcb4 undergoing spontaneous open-and-close motions. By aligning the twelve structures based on the first lobe, we found that the motion of the second lobe follows a defined trajectory, in which the two lobes of DrAbcb4 not only open progressively wider laterally as measured by D_COM_ but also undergo angular rotations, as demonstrated for the NBD2 (Fig. 2A). Such an opening-while-twisting motion can be directly visualized in the 3D Variability Analysis in cryoSPARC (Supplementary Movies 1 and 2), from which continuous conformational changes could be resolved^31^. The spontaneous open-and-close motions of DrAbcb4 may present an important feature of the transporter actively sampling for potential ligand at resting state but may also form the basis for the basal ATPase activity.

**Figure 2.**
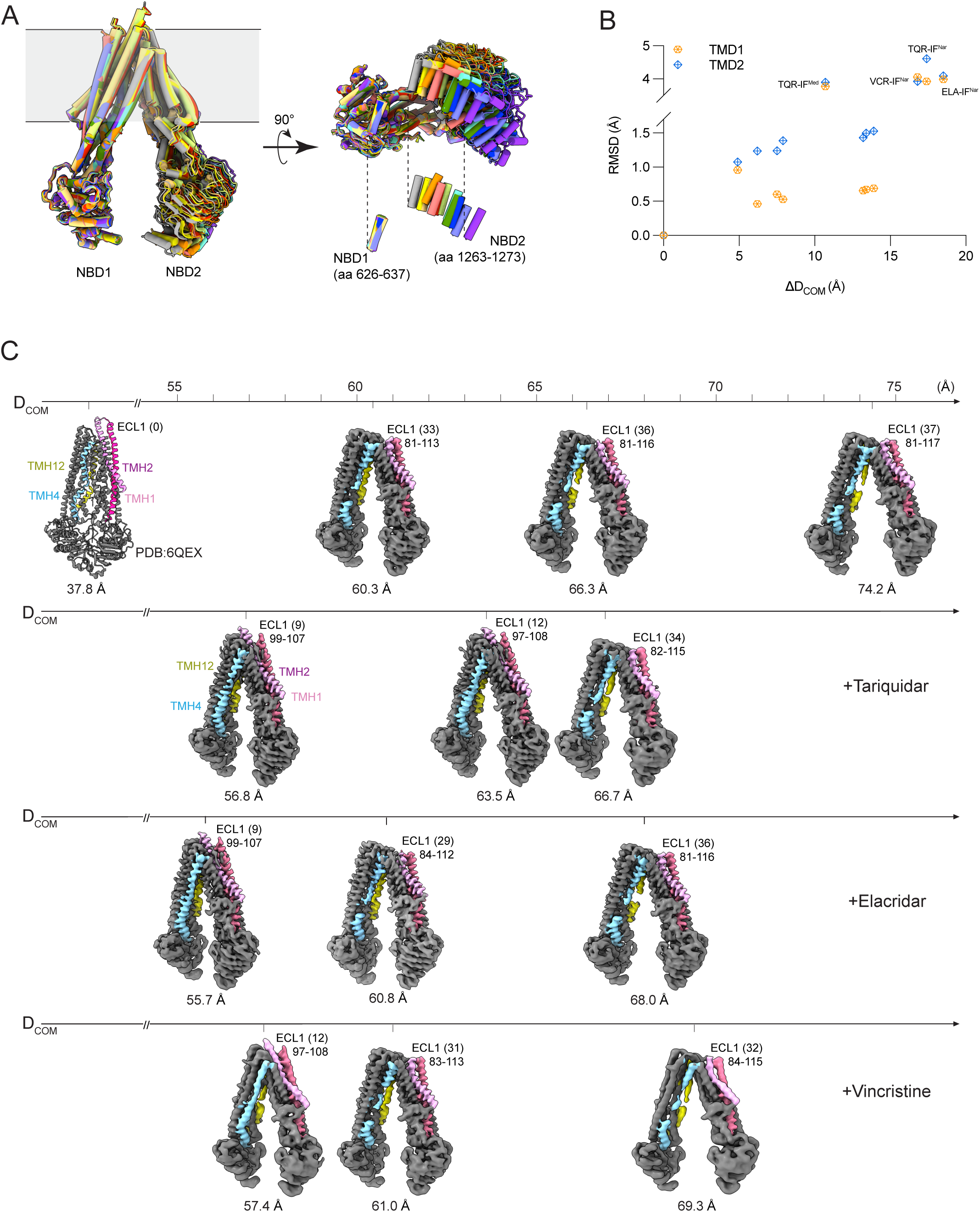
Coupled structural changes in DrAbcb4 as a function of open-and-close movement. (A) The two lobes of Abcb4 follows a defined trajectory while undergoing the open-and-close movement. Twelve structures of DrAbcb4 with different D_COM_ are aligned based on their N-terminal lobe, and the trajectory is indicated by the positions of a helix (residues 1263-1273) in NBD2 of the C-terminal lobe. (B) Using Apo IF^Wide^ as a reference for superposing the 12 Abcb4 structures pairwise, the rms deviations (RMSD) are plotted against the changes in D_COM_ (ΔD_COM_), revealing a positive correlation indicative of coordinated movement. (C) Structural changes in TM helices in relation to the open-and- close motion. Structure of hP-gp in narrowly open IF occluded conformation (PDB:6QEX) together with 12 high-resolution EM density maps of Abcb4 from the four datasets are shown. Helices TMH1 and TMH2 are colored pink and magenta, respectively. TMH4 is colored cyan and TMH12 is in yellow.

The 3D Variability Analysis also shows that, rather than moving as rigid bodies connected by a flexible linker, each lobe of DrAbcb4 undergoes significant local conformational changes, especially for TM helices in the TMDs, during the open-and-close motions. These local changes can be detected by structure superpositions. By pairwise aligning TMD1 or TMD2 of DrAbcb4, using IF^Wide^ (D_COM_=74.2Å) as reference, we observed that the structural difference represented by RMSD is a function of the differences in D_COM_ (ΔD_COM_) (Fig. 2B). Specifically, models with similar D_COM_ values exhibit lower RMSD for both TMDs, whereas models with very different D_COM_ show higher RMSDs. Notably, all four substrate- or inhibitor-bound structures exhibit disproportionally large RMSDs, reflecting their tendency to maintain the conformation of small D_COM_. This observation also extends to the superposition that included the structure of mP-gp (PDB:5KPD, D_COM_=47.2Å) as reference (Supplementary Figure 8). Thus, we conclude that the structures of mP-gp and DrAbcb4 are comparable.

Notably, as the two lobes open wider and D_COM_ increases, (1) TMH1 and TMH2 unwind at their tips on the extracellular side of the membrane, extending the unstructured ECL1 loop (Figs. 2C and 3A, Supplementary movie 2). The ECL1 loop in hP-gp forms part of the epitope recognized by the conformation-sensitive monoclonal antibodies MRK16 and UIC2, indicating that these antibodies are likely responsive to different conformations of hP-gp. (2) Both coupling helices TMH10 and TMH11 of TMD1 undergo a large positional shift (Fig. 3B). (3) By contrast, only TMH4 of the coupling helix pair TMH4 and TMH5 of TMD2 undergoes similar positional changes (Fig. 3C). (4) Furthermore, the mid-section of TMH4 unwinds, leading to its breaking up gradually at two mid-sections (residues S233-G238, and T245-T250), forming three noncontinuous shorter helices. Similarly, the density for the mid-section (residues F992-Y996) of TMH12 progressively deteriorates, unwinding into a loop that connects the top and bottom halves of the TMH12 (Fig. 2C). Unwinding of a stable helix is considered energetically unfavorable in protein folding^32^, but it could be an important investment to have sufficient clearance to bind larger substrates. The energy source that drives the unwinding is still undetermined and coupling of the open-and-close motions to individual helix unwinding may be part of the solution.

**Figure 3.**
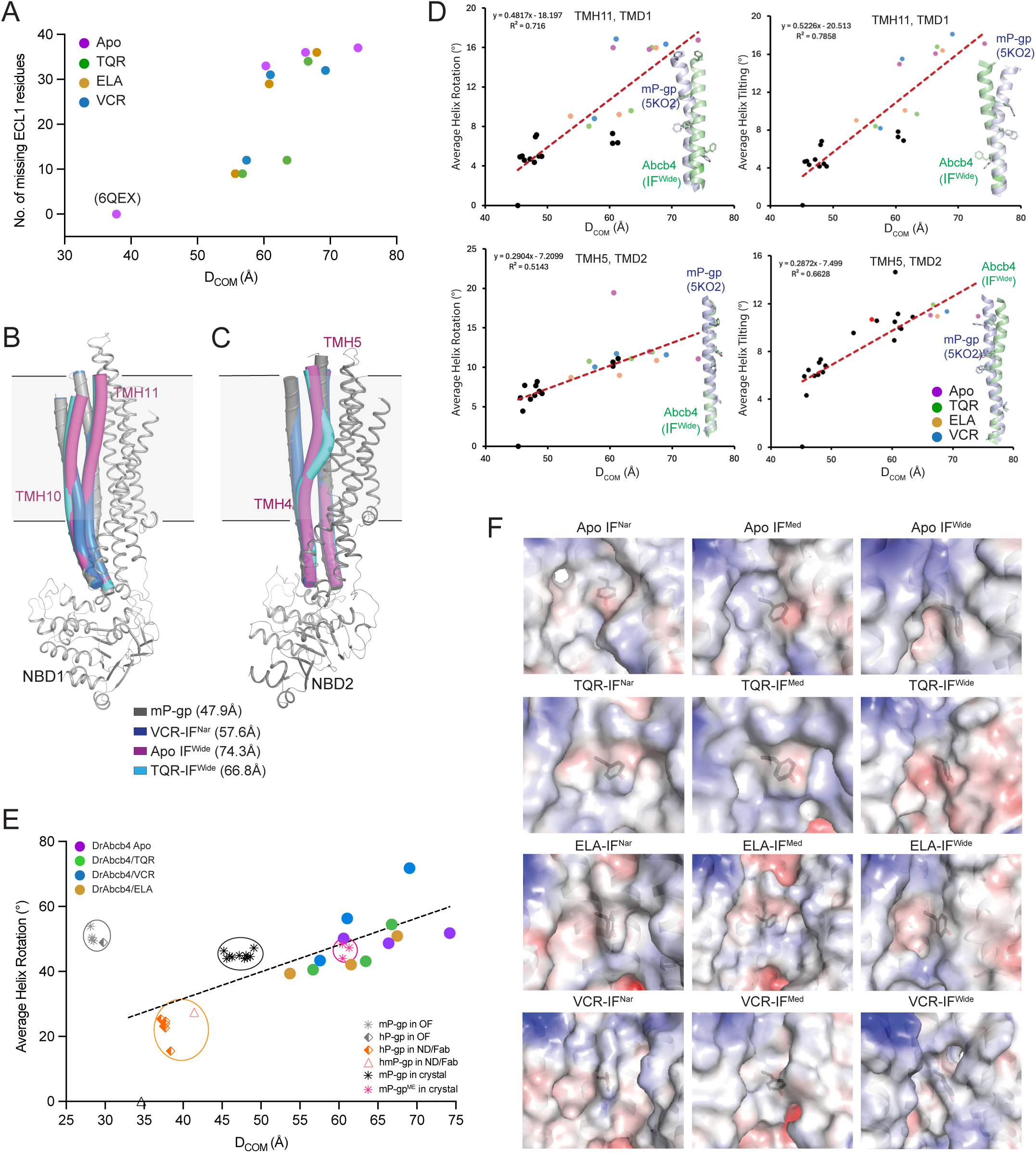
Coupling between conformational changes in TM helices and open-and- close movement. (A) The number of disordered ECL1 residues as a function of D_COM_. (B) Positional shifting of coupling helices TMH10 and TMH11 of TMD1 in response to changing D_COM_. N-terminal halves of four structures are aligned and rendered: mP-gp (5KDP), VCR-IF^Nar^, Apo-IF^Wide^, and TQR-IF^Wide^. Only the mP-gp structure is presented in full as ribbon diagram. The others are shown with TMH10 and TMH11 only as tubes. The extracellular portion of the TMH10 and TMH11 have their positions shifted for Apo-IF^Wide^ and TQR-IF^Wide^ compared to those of mP-gp and VCR-IF^Nar^. (C) Positional shifting of TMH4 of TMD2 in response to changing D_COM_, shown similarly as in (B). The extracellular portion of the TMH4 has its position shifted in Apo-IF^Wide^ and TQR-IF^Wide^ compared to those of mP-gp and VCR-IF^NAR^. (D) Scatter plots showing the correlation of D_COM_ with average rotation angle <HRA> and average tilting angle <HTA> for the two coupling TM helices, TMH11 for TMD1 and TMH5 for TMD2. Black dots are from crystal structures of mP-gp or methylated mP-gp, colored dots are for Abcb4 structures. The dashed trending lines through sample dots are given in red with fitting equations and R values. The two helices in cartoon rendition in each panel are from mP-gp (purple) and Abcb4 Apo IF^Wide^ (green), respectively, showing in one of the two different orientations in each panel for a given helix. Equivalent aromatic residues are shown as stick models. (E) Scatter plot of <HRA> of TMH10 as a function of D_COM_ showing clustering of P-gp structures obtained under various conditions. The dashed line is trending through the sample dots minus those in the OF conformation. (F) Changes in the substrate-binding environment shown as electrostatic surface potential centered on the conserved residue Y951 (in stick model) of TMD1 in Abcb4 structures of different D_COM_s. Positive potential is colored in blue, negative in red, and neutral in white.

### Conformational plasticity of TM helices in inward-facing conformations

The conformational plasticity of TM helices observed in the structures of DrAbcb4 accompanied by varying D_COM_ values was reminiscent of our previously reported work describing crystal structures of mP-gp in the absence of bound drugs^27^, in which Esser et al. also observed coupled movements between D_COM_ and rotation and tilting of individual helices. The coupling of each individual helix motion to the open-and-close movement is complex; nevertheless it can be described partially by measuring for each helix the average helix rotation angle (<HRA>, in degree) and the average helix tilting angle (<HTA>, in degree) and plotting against D_COM_, as reported before^27^. To cover a larger D_COM_ range, we also included in this plot crystal structures of mP-gp and methylated mP- gp (mP-gp^ME^) (Fig. 3D, Table 1, and Supplementary Figure S9). Clearly, there is a strong correlation between the D_COM_ and the average helix rotation/tilting angles with the correlation coefficient in the range of 0.6-0.9. We noticed that in drug bound structures such as DrAcb4/TQR, some helices display large <HRA> and <HTA> values, deviating significantly from trending lines, indicating involvement of residues in these helices in drug binding.

Plotting <HRA> and D_COM_s for P-gp structures available in the PDB showed interesting clustering of structures (Fig. 3E). Some of the structures are off the trend line by large margins, lowering the correlation coefficient significantly if included in the calculation. The clustering correlates different conditions under which these structures were determined. For example, all structures from crystallized mP-gp are clustered together, showing very narrow D_COM_ and <HRA> ranges, and so are cryo-EM structures of hP-gp imaged with bound monoclonal antibody and embedded in nanodiscs, as well as those in the outward-facing (OF) conformation, indicating conformation restrictions due to experimental conditions or to functional states.

The movement of individual helices, coupled with the open-and-close motion of the two lobes, should conceivably lead to a dynamically shifting surface topography of the drug-binding site to enable binding to different drugs. To visualize the dynamics of the drug-binding site, we chose the residue Y951, located in TMD1 of DrAbcb4, which is completely conserved in murine and human P-gp and involved in contacts with nearly all compounds bound to P-gp (Supplementary Figure 10). The environment centered on Y951 is rendered as electrostatic surface potential for each DrAbcb4 structure (Fig. 3F). Clearly, the surface variations among DrAbcb4 structures with different D_COM_ values support our finding that correlated movement of individual helices and the open-and-close motions of the two lobes potentially allow DrAbcb4 to adopt many electrochemical and topographical environments within the drug-binding pocket to interact with drugs of diverse sizes and constituents.

### Versatile binding of drugs to DrAbcb4

Conceivably, the fluid nature of the drug-binding surface should also enable a substrate to bind at multiple sites and/or in different conformations during protein-drug interaction. This hypothesis gained support from various inhibitor/substrate conformations or poses identified in our DrAbcb4 structures with bound drugs. In the final high-resolution DrAbcb4 maps obtained in the presence of drugs, additional EM densities, at similar contour levels to those of proteins, were observed between the two TMDs after completion of protein model building. The interpretability of these densities improved following maximum likelihood-based refinement using the two half maps^33^.

Among the three DrAbcb4/TQR maps with different D_COM_ separations, only the TQR-IF^Nar^ (D_COM_=56.8Å) and TQR-IF^Med^ (D_COM_=63.5Å) were found with bound tariquidar molecules. For the TQR-IF^Nar^ map, two tariquidar molecules were built into continuous density located at the central drug-binding cavity in a back-to-back manner, touching each other briefly (Fig. 4). As it turns out, this back-to-back arrangement of bound molecules appears to be a common feature, although the two molecules need not be identical. Of the two modeled tariquidar molecules, designated as TQR-1 and TQR-2, TQR-1 is facing the TMD1 side of the drug-binding cavity, whereas TQR-2 resides in the side of TMD2 (Fig. 4C and Supplementary Figure 10). Under this specific DrAbcb4 conformation, TQR-1 has contacts with 10 residues at a distance cutoff of 3.5Å, all of them hydrophobic with four aromatic ones. The two methoxy groups on the isoquinoline moiety of TQR-1 are about 3.6Å from the OH group of Y951, forming potential hydrogen bonds. The quinoline carboxymide group at the other end of TQR-1 is anchored by two aromatic residues (F77 and Y981) (Fig. 4D). TQR-2 interacts with 13 residues; ten are hydrophobic including five aromatic residues, and three are hydrophilic. Its quinoline moiety wedges into the gap between the pair of coupling TMHs 4 and 5, contacting A234, A235, and K239 on TMH4 and residue T307 on THM5 (Figure 4D). TQR-2 likely forms three hydrogen bonds with Y315, T307, and K239. One must note that TQR-1 and TQR-2 have very different conformations.

**Figure 4.**
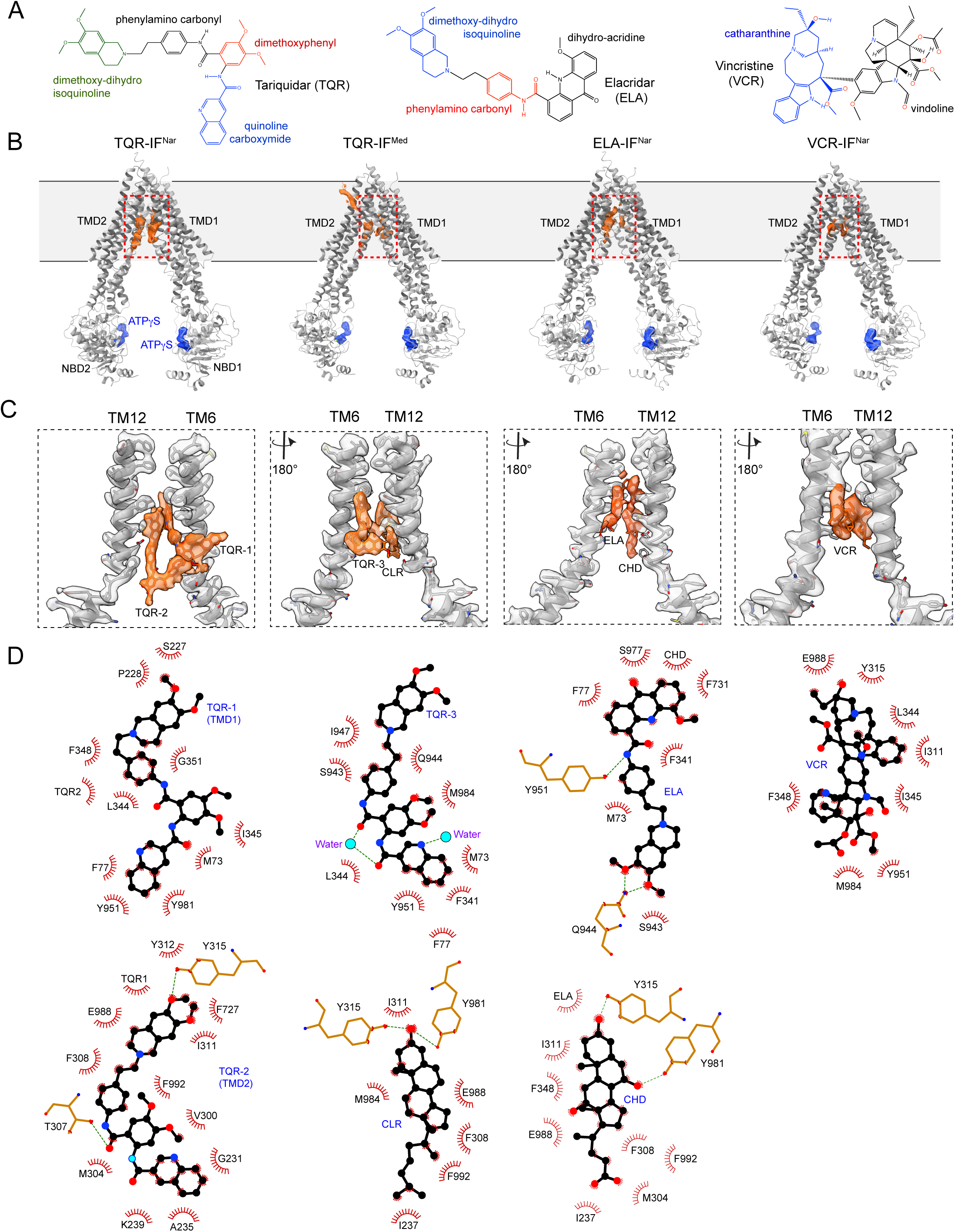
Versatility in drug binding. (A) Chemical structures of tariquidar, elacridar, and vincristine. (B) Bound drugs revealed by Abcb4 structures. Four Abcb4 structures in ribbon diagram are presented: from left to right TQR-IF^Nar^ with two bound tariquidar (TQR) molecules, TQR-IF^Med^ with one bound tariquidar and one cholesterol (CLR), ELA-IF^Nar^ with one bound elacridar (ELA) and one cholate (CHD), and VCR-IF^Nar^ with one bound vincristine (VCR). The EM densities for bound drugs are shown in orange, and densities for ATPγS/Mg^2+^ are shown in blue. The two parallel lines depicting the membrane bilayer. (C) Zoom-in views of the drug-binding cavities highlighted by red rectangles in (B). Overlapping EM density with ribbon structures for TMHs 6 and 12 (gray) as well as bound drugs (orange) are shown. (D) Ligplot diagram depicting conformations of bound drugs and their protein environments. Bound drugs are shown as ball-and-stick models with carbon atom in black, nitrogen atom in blue and oxygen in red. Residues that are in close contacts with the bound drugs are given as eyelash icons in red, and those may form H-bond are given in stick models.

For the TQR-IF^Med^ map, only one tariquidar (TQR-3) was modeled into the EM density at the TMD1 side of the binding cavity (Fig. 4B and 4C). In contrast to TQR-1 and TQR-2 that display extended conformations, TQR-3 assumes a more compact pose, being embraced by nine mostly hydrophobic residues (Figure 4D). There is a cholesterol molecule (CLR) identified in the TMD2 side of the cavity, juxtaposing TQR-3 and contacting its dimethoxy-isoquinoline moiety. These results demonstrate that DrAbcb4 is capable of binding compounds in different protein conformations (different D_COM_ values), using various available sites of the binding cavity, and even handling many conformers of drugs.

Similar observations were made for reconstructions of DrAbcb4/elacridar and DrAbcb4/vincristine. The data set of the DrAbcb4/elacridar sample produced three reconstructions, of which only the ELA-IF^Nar^ map revealed the inhibitor bound at the TMD1 side of the drug-binding cavity, which is supported by an additional density that was best fit with a cholate molecule used in sample preparation (Fig. 4D). The bound elacridar is surrounded by eight residues, mostly hydrophobic, and its terminal dihydroacridine moiety is embedded in six aromatic residues.

One vincristine molecule was found to occupy the entire cavity in the VCR-IF^Nar^ structure. Due to its large size (825 Daltons), vincristine interacts with a total of eight residues from both TMDs, mostly hydrophobic (Fig. 4). Specifically, its larger vindoline moiety occupies the TMD1 side of the cavity, whereas the smaller catharanthine group interacts with TMD2 part of the binding site. Unlike the structures of TQR-IF^Med^ and ELA-IF^Nar^ that have juxtaposing cofactors, VCR-IF^Nar^ features no extra density and no additional support for the binding of vincristine. Notably, some bound drugs may be modeled in two different poses revealed by difference density maps, as branching density is observed in the reconstruction, likely resulting from a mixture of multiple transient conformations with small differences inseparable by 3D classification (Data not shown).

### Structural conservation of the drug-binding site in DrAbcb4

To understand the significant conservation in substrate specificity profile between DrAbcb4 and P-gp, we examined sequence conservation for amino acid residues involved in drug binding. We first surveyed 15 published structures of P-gp, including hP-gp and mP-gp bound with various compounds^17–21,30^. These represent a total of four different P-gp IF conformations and 59 drug-interacting residues (Supplementary Figure 10). The latter exhibits a sequence identity and similarity of 73% and 97%, respectively, exceeding those of the overall values of 66% and 80%, respectively.

We next asked how our limited number of DrAbcb4 structures with bound inhibitors expands the repertoire of residues involved in substrate binding. With five bound compounds in DrAbcb4 structures in four different conformations, we found a total of 31 drug-interacting residues. Among these, 23 residues overlapped with those found in P-gp, while eight additional residues were identified in this work, bringing the total number of drug-interacting residues to 67 (Supplementary Figure 10). The sequence conservation for the drug-binding residues remains unchanged.

Overall, TMD2 contributes more to drug binding, as two-thirds of the 67 drug-binding residues are from TMD2. Among them, the conserved residue Y315 in TMH5 of DrAbcb4 (Y306 in mP-gp and Y310 in hP-gp) is on top of the list, interacting with various compounds in 18 out of the 19 surveyed structures. It is followed by Y981 in TMH12 (F979 in mP-gp and F983 in hP-gp) and F348 in TMH6 (F339 in mP-gp and F343 in hP-gp), which appeared 16 and 15 times, respectively. The wider separation distances observed in DrAbcb4 have made available more residues for drug binding. The highly conserved drug-binding pocket likely underlies the largely overlapping substrate/inhibitor specificity profiles of P-gp and DrAbcb4. Therefore, zebrafish Abcb4 is not only a functional homolog of human P-gp but also shows significant structural similarity, particularly in drug-binding regions, making it an appropriate model system for structural and mechanistic studies of P-gp.

## Discussion

Modeling P-gp function at various physiological barriers and understanding the mechanism of its substrate polyspecificity have been subjects of active research. Just as for P-gp in mammals, DrAbcb4 contributes to xenobiotic resistance of zebrafish embryos via its efflux activity. It was shown to be the functional equivalent of P-gp with an overlapping substrate specificity and tissue localization profile^24–26^. In this study, the similarity extends beyond function and structure to include the mechanism of function, as revealed by the structures of DrAbcb4, both drug-free and drug-bound, in the presence of ATPγS/Mg^2+^. Abcb4 remains in IF conformations with a range of separations between its two lobes and with well-defined ATPγS/Mg^2+^ density in both NBDs (Fig. 1 and Supplementary Figure 3D). Notably, these structures were obtained without the use of mutations, conformation-selective antibodies, crosslinking, nanodisc embedding, or crystal contacts during sample preparation. The structures illustrate the extent that an unrestricted Abcb4 could achieve in the separation of the two lobes and reveals that drug binding, rather than nucleotide binding, promotes the two lobes to come closer. Moreover, the coupling of the open-and-close motions of the two lobes to movements of individual TMHs is consistent with the multidrug transport function of DrAbcb4. The structural similarity between P-gp and DrAbcb4, especially at the substrate-binding site, explains their shared substrate/inhibitor-specificity profile and indicates a homologous transport mechanism.

In addition to the structural, functional, and mechanistic similarities of DrAbcb4 and hP-gp, the use of DrAbcb4 in these studies has several other advantages. DrAbcb4 can be genetically deleted and mutated within zebrafish^34^, where it plays a critical role at the blood-brain barrier and determines the uptake of many drugs in the intestine and their excretion by the kidney and liver^22,24–26^. Observations made concerning residues involved in drug binding, in proposed mechanisms of polyspecificity and P-gp function, and in searching inhibitory agents can be readily tested in a moderately high throughput animal model. Furthermore, DrAbcb4 has been easy to express in a *Pichia*-based expression system, yielding large amounts of homogeneous protein and showing favorable solution properties that are quite suitable for cryo-EM analysis.

### Conformational landscape revealed by Cryo-EM structures of DrAbcb4

Previously reported IF structures of P-gp showed the NBD separation distances (D_COM_) clustered between 38Å and 60Å (Table 1 and Fig. 1). For example, all crystal structures of mP-gp have their D_COM_ clustered around 46Å^15,27^; all crystal structures of methylated mP-gp cluster around 61Å^17,27^; and all Fab-bound hP-gp cluster around 38Å^18–20^, indicating that these experimental conditions likely restrict movement of the two lobes. By contrast, the DrAbcb4 structures show no such clustering with D_COM_ values distributed over the range between 45Å and 74Å (Fig. 1B). Cryo-EM datasets also survey all particles in solution (on grid) that minimize any uncontrolled experimental biases. Despite concerns over the presence of detergent in sample preparation, multiple lines of evidence suggest that the wider separations of the two lobes observed in unrestricted DrAbcb4 structures may be functionally relevant. First, restricting the motion of P-gp through linker manipulations did not alter its structure but abolished its drug transport activity and drug-stimulated ATPase activity^27,28^, suggesting that linker flexibility enabling P-gp to open wider is essential for its function. Second, supporting this hypothesis, a recent study on the bacterial ABC transporter MsbA, whose wide-open IF structure was once thought to be an artifact, found this conformation to be predominant among various IF conformations in live *E. coli* cells, likely allowing large substrates to access the binding cavity^35^. Thirdly, double electron-electron resonance (DEER) studies on P-gp observed similar wide-open IF conformation (>70-80 Å) in its apo, substrate-bound, or nucleotide-bound states^36^. Fourth, atomic force microscopy (AFM) study, in real time, of hP-gp embeded in the membrane bilayer revealed a 28% probability of its two lobes existing in the "extreme IF" conformation^37^. Therefore, the open-and-close motion may represent a common mechanism employed by P-gp-like ABC transporters to capture substrates from the membrane.

Additionally, Loo and Clark used double cysteine mutations to crosslink TM6 (L339C) and TM12 (V982C). The amount of crosslinked product was decreased in the presence of ATP. Another pair of cysteine mutation crosslinks TM6 (F343C) to TM12 (V982C) does the opposite, with increased crosslink product in the presence of ATP^38^. The authors suggested that helix rotation, either a single helix or both, must be involved.

Examination of the hP-gp, mP-gp, and DrAbcb4 structures with varying D_COM_ separations reveals that, in addition to helix rotation, crosslinking between the L339C and V982C pair can occur only when the two lobes of P-gp have a D_COM_ greater than 65Å.

### Coupling of the open-and-close motion to individual TM helix movement enables substrate polyspecificity

Although relative rearrangements of TM helices have been reported in ABC transporter structures, especially during the IF to OF transition, our finding that individual TM helices within TMDs move relative to each other during the IF open-and-close motions challenges the conventional depiction of TMDs moving as rigid bodies. Moreover, the complex maneuvers of these helices, including rotation, tilting, twisting, and positional shifting, are shown to couple to the open-and-close motion of the two lobes, which is evident from the 3D Variability Analysis of EM data (Supplemental movies 1 & 2) and from the correlative analysis of helix rotation and tilting with D_COM_ (Fig. 3D and Supplementary Figure 9). This correlative analysis is also able to cluster together structures that are obtained under specific restrictive conditions, such as packed within the crystal lattice or inhibited by Fab binding, illustrating that not only do these structures have a small range of D_COM_s but also give a similar average of helix rotation and tilting angles (Fig. 3E). The coupled open-and-close motions and individual TMHs movement observed in our cryo-EM analysis inevitably leads to a fluid substrate-binding surface that is continuously changing in shape, size, and chemical composition during the transport cycle (Fig. 3F); this also explains why crosslinking a substrate to a specific TMH residue is able to arrest P-gp in a single conformation^21^.

The relative rotation of individual TM helices during hP-gp’s functional cycle was previously suggested by Loo and Clark using crosslinking experiments^38^ and its coupling to the global open-and-close motions was later reported based on structural studies of mP-gp^27^. This coupling led to the hypothesis that a single substrate may bind to P-gp at multiple sites of the binding cavity involving different residues and the substrate can assume different conformational poses, which could underlie the structural basis for the polyspecificity of P-gp^39^. This dynamic nature of both the protein’s binding surface and the substrate’s conformation stands in sharp contrast to the classic lock-and-key model traditionally used to explain enzyme-substate specificity. Instead of a rigid binding site, the fluid binding surface in P-gp’s drug-binding cavity could enable binding of a diverse range of substrates. Ligands, in turn, may also adopt different poses to match the changing pocket. Indeed, this synergistic relationship is exemplified by the distinct conformations adopted by tariquidar when bound to DrAbcb4 or P-gp in different IF states (Fig.4 and Supplementary Figure 11). Strikingly, none of the five bound tariquidar molecules assumes the same pose. Similar observations were made for elacridar and taxol (Fig. 4 and Supplementary Figure 11). Supporting this, previous photoaffinity labeling experiments with P-gp using propafenone-type compounds revealed multiple covalently attached drugs to at least four TM helices^40^.

### Proposed mechanism of substrate polyspecificity and modulated ATPase activity

The conformational dynamics and landscape provided by the structures of DrAbcb4 and P-gp, along with coupled movement of TM helices, explains P-gp’s broad substrate specificity. In the absence of drugs, the two halves of P-gp undergo continuous open-and-close motions, which allows the drug-binding cavity to sample different states in anticipation for potential drug entry. As the transporter cycles through the open-and-close motions going back and forth between IF^Wide^ to IF^Nar^ states, occasionally, the two lobes transition to the state previously described as "IF occluded state" where the binding cavity is closed to both sides of the membrane^19^, before eventually proceeding to active ATP hydrolysis. This process is relatively slow, with an approximate turnover rate of one ATP per second, which is known as the basal ATPase activity (Fig. 5A).

**Figure 5.**
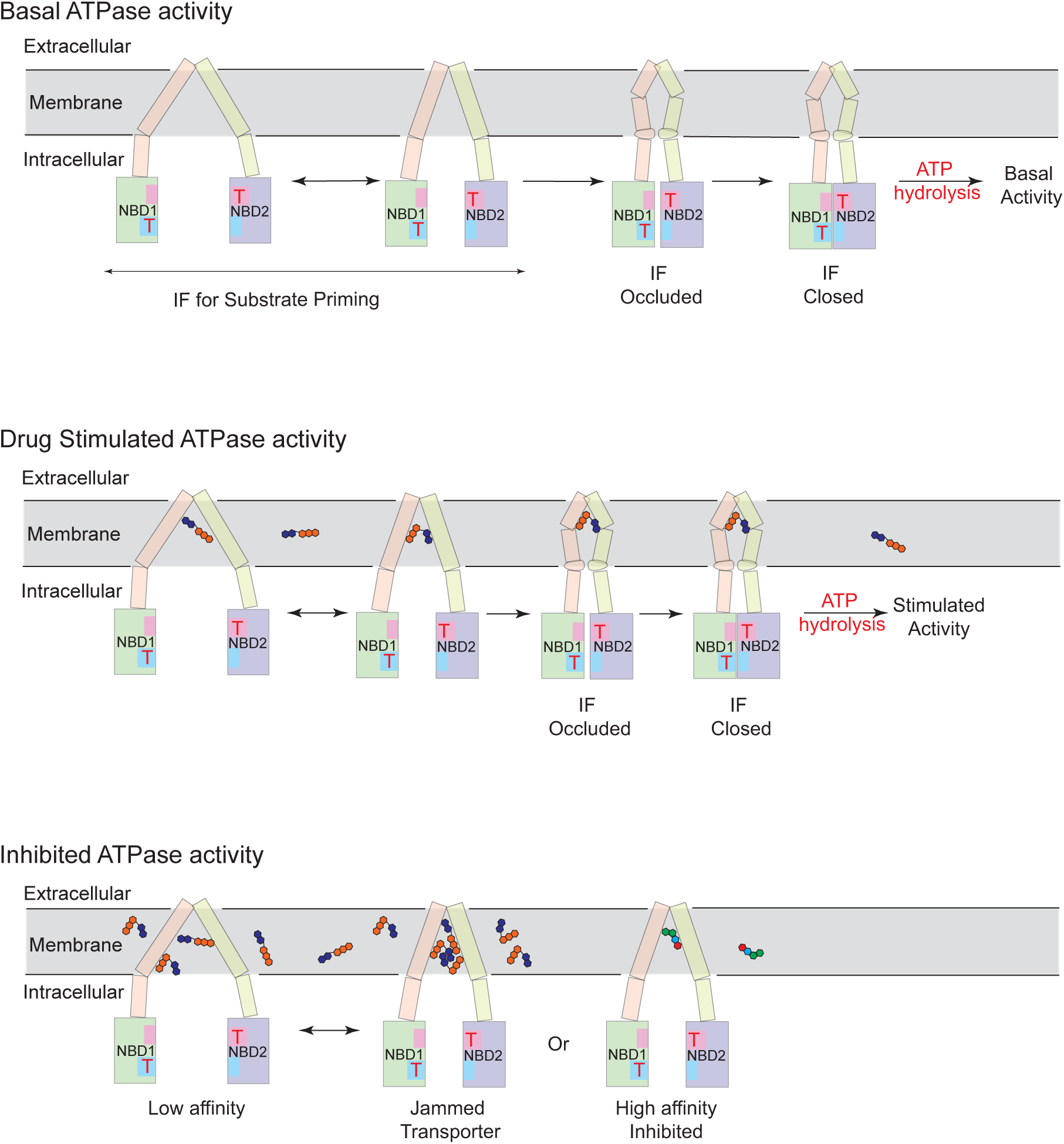
Proposed mechanism of substrate recognition and ATPase activity modulation in P-gp. P-gp or Abcb4 in membrane undergoes spontaneous open-and-close motion of its two lobes. The two homologous NBDs are green and purple rectangles and bound ATP are labeled as Ts in red. The two TMDs are represented by sticks in pink and yellow, respectively. The drug-binding cavity encompassed by the entire TMD can be used for substrate interactions. The "IF occluded state" is characterized by the substrate-binding cavity closed (not the NBDs), which proceeds to ATP hydrolysis via the "IF closed state" (NBDs closed). In the absence of substrate, the transitions from open-and-close motion to "IF occluded state" and to the "NBD closed state" may be slow leading to basal ATPase activity (top panel). In the presence of substrates represented by five connected circles, both protein and drug undergo conformational changes, leading to accelerated transitions to the "IF occluded state" and to the "NBD closed state", followed by stimulated ATP hydrolysis (middle panel). Inhibition of P-gp may occur by halting the open-and-close movement of its two lobes or any of the conformational transitions, either by jamming or by high affinity specific binding (bottom panel).

Following this notion, some ligands could modulate the ATPase activity by either accelerating or slowing down the conformational transitions that ultimately leads to ATP hydrolysis. Substrates enter the binding pocket through the gap between TMD1 and TMD2 and subsequently bring the two lobes together, facilitating a more rapid transition to the “IF occluded state", which leads to a stimulated ATPase activity (Fig. 5B). A key aspect of this model is that P-gp can handle a vast range of substrates. Drug binding can occur across various IF conformations, with single or multiple drugs bound simultaneously in different poses^19–21,41^. One question arises as to how P-gp prevents the bound substrates from escaping its binding cavity? A likely solution is through the previously described "IF occluded state" (Fig. 5)^19^, which can be obtained in the absence of stabilizing monoclonal antibody^30^ ^42^. Conceivably, as the two lobes clamp down, bound compounds are exposed to more residues and form more interactions with residues from both lobes, while the compounds are adjusting their poses accordingly (Fig. 4 and Supplementary Figure 11), until the "IF occluded state" is reached.

On the other hand, the ATPase activity could be inhibited by a reduction in the motion of the two lobes of P-gp. Some compounds like tariquidar may bind P-gp with higher affinity in certain IF conformations, likely halting the open-and-close movement and slowing down ATPase activity (Fig. 5C), which is supported by a reduction in the basal activity^19^. Thus, a reduction in the motion of the two lobes of P-gp may represent one of the inhibitory mechanisms. Similar mechanism could be applied to inhibiting P-gp by cross-linking TM helices with drugs^21^. Compounds with relatively weak affinity that typically accelerate ATPase activity at low concentrations may also inhibit the conformational transition by jamming the transporter if present at high concentrations.

## Materials and Methods

### Expression of DrAbcb4 in *Pichia pastoris*

The expression of DrAbcb4 in *Pichia pastoris* followed the protocol described in EasySelect Pichia Expression Kit (Life Technologies, Thermo Scientific, MA). The gene DrABCB4 encoding protein (UniProt accession number E7F1E3) was subcloned from a plasmid with full-length DrAbcb4 in pcDNA3.1 vector^26^ into a Pichia yeast expression vector pPICZ-A (Life Technologies) with a poly-histidine tag at the C-terminus (pPICZ-DrAbcb4-hexahis). Sequences of the plasmids were confirmed before transfection.

The plasmid pPICZ-DrAbcb4-hexahis was linearized using the restriction enzyme PmeI (New England BioLabs) before being introduced into *Pichia pastoris* yeast strain KM71H via electroporation. Transformants were initially grown at 30°C in a buffered minimal glycerol medium (100 mM potassium phosphate, pH 6.0, 1.34% YNB, 4 x 10^-5^ % biotin and 1% glycerol) until optical density at 600 nm (OD_600_) reached 6. Cells were then pelleted down and resuspended in a buffered minimal methanol medium (100 mM potassium phosphate, pH 6.0, 1.34% YNB, 4 x 10^-5^% biotin and 0.05% methanol) with the OD600 reading close to 1. The resuspended cells were grown at 30°C for 24h before centrifugation at 4,000 x g for 25 min to harvest the cells. The cell pellet was kept at - 80°C until use.

### Isolation of the membrane fraction from *Pichia pastoris*

Frozen cell pellets were thawed at 4°C and resuspended in homogenization buffer (100 mM Tris, 100 mM Sucrose, 100 mM 6-aminohexanoic acid, 1 mM EDTA, 2 mM PMSF, 2 mM DTT, pH 7.5) using a handheld Dounce homogenizer. The resuspended cells were applied onto a Microfluidizer (Microfluidics International Cooperation) at 900 bars for three cycles to lyse the cells. The insoluble cell debris were separated from the supernatant by centrifugation at 3,500 x g for 20 min. To isolate the crude membrane in the supernatant, multiple ultracentrifugation steps at 100,000 x g were performed and each time only the pellet was kept. After the first 90-min ultracentrifugation, the pellet was homogenized in the same homogenization buffer and pelleted down again by a second ultracentrifugation at 100,000 x g for 90 min. Then, the pellet was homogenized in wash buffer (25 mM Tris, 200 mM NaCl and 5 mM β-mercaptoethanol, pH 7.5) followed by a third ultracentrifugation at 100,000 x g for 30 min. The pellet after the 3^rd^ ultracentrifugation was homogenized in a storage buffer (25 mM Tris, 30 % glycerol, 5 mM β-mercaptoethanol, pH7.5) and kept at -80°C until use.

### Purification of DrAbcb4 from isolated membrane

Frozen membrane was thawed at 4°C. After determining the total protein concentration in the membrane using the Pierce BCA Protein Assay Kit (Thermo Fisher Scientific), the protein concentration was adjusted to 5 mg/ml using the solubilization buffer (10 mM Tris, 30 mM Imidazole, 75 mM NaCl, 15% glycerol, 5 mM β-mercaptoethanol, pH7.5). While stirring on ice, 20% DDM (Anatrace) stock solution was added slowly to a final concentration of 2% to solubilize the membrane. After 1 hour of stirring, the solubilized membrane was subjected to ultracentrifugation at 100,000 x g for 30 min. In the meantime, the Ni-NTA resin (Qiagen) was treated with buffer A (20 mM Tris, 75 mM NaCl, 15% glycerol, 2 mM β-mercaptoethanol, 0.0675% DDM, 0.04% Na cholate, pH7.5) supplemented with 30 mM Imidazole. The supernatant after the ultracentrifugation was mixed with pre-equilibrated Ni-NTA resin at 4°C for 2 hours before being packed into a gravity flow column. The unbound proteins were washed away using 10 column volumes of buffer A supplemented with 30 mM Imidazole, and the bound Abcb4 protein was eluted with buffer A supplemented with 300 mM Imidazole. The eluate was further concentrated using a 100 kDa cut-off Amicon centrifugal filter (Fisher Scientific) and loaded onto a Superdex^TM^ 200 Increase 10/300 GL column (Cytiva) pre-equilibrated with SEC buffer (20 mM Tris, 75 mM NaCl, 2% glycerol, 2 mM β-mercaptoethanol, 0.0675% DDM, 0.04% Na cholate, pH 7.5). The flow rate was kept at 0.5 ml/min, and the fractions were collected.

### ATPase activity

The ATPase activity of zebrafish Abcb4 was determined by quantitating the green complex formed between malachite green, molybdate, and the inorganic phosphate released from ATP hydrolysis^43,44^ of Abcb4. A total of 50 μl reaction containing 4-8 ng of protein in the assay buffer (50 mM HEPES, 10 mM MgCl_2_, 4 mM ATP and 5 mM DTT, pH 7.5) was incubated at 30°C. After 40 minutes, the reaction was stopped immediately by adding 800 μl of STOP solution (a fresh mixture of 0.045 % malachite green and 1.4 % ammonium molybdate tetrahydrate in 4 N HCl in a 1:3 ratio). After 60 seconds, 100 μl of 34% sodium citrate was added and mixed. The admixture was incubated in room temperature for 10 minutes, then 16 μl of 20% Tween-20 was added to dissolve any precipitations. A standard curve was established using a series of known amount of KH_2_PO_4_ (10-100 μM) dissolved in the assay buffer following the same procedure. Absorbance at 660 nm was measured for all the samples, and the amount of inorganic phosphate released was calculated based on the standard curve.

To measure the ATPase activity of Abcb4 reconstituted into liposome, total soy lipid extract was dissolved in 50 mM HEPES buffer, pH7.5, to a concentration of 20 mg/ml and sonicated gently until clear. Abcb4 was incubated with lipid solution at 1:1 ratio (w/w) on ice for 1 hour before the assay. Verapamil stimulated ATPase activity was measured by pre-incubating Abcb4 in the same assay buffer supplemented with a series of concentrations of verapamil at 30 ℃ for 5 minutes before adding ATP to start the reaction.

### Cryo-EM grid preparation and data collection for DrAbcb4

The peak fraction of DrAbcb4 (6 mg/ml), collected from size exclusion chromatography, was incubated with 10 mM of ATPγS and 10 mM of MgCl_2_ on ice for about 30 minutes before freezing grid for the drug-free Abcb4 sample. For drug-bound samples, stock solutions of the compounds dissolved in DMSO were added to the final concentrations of 0.28 mM for tariquidar, 0.24 mM for elacridar, or 0.45 mM for vincristine, along with 5 mM ATPγS and 5 mM MgCl_2_. These samples were incubated on ice for 2 hours, followed by a 30-minute centrifugation at 20,817xg at 4°C to remove any precipitates prior to grid freezing.

For grid preparation, Quantifoil R 0.6/1 200-mesh Cu grids or UltrAuFoil R 0.6/1 300-mesh grids were glow-discharged using a PELCO easiGlow for 60 seconds with a current of 15 mA. Using a Vitrobot Mark IV (ThermoFisher Scientific) set at 95% humidity and 4°C, 2.5 ul of sample was applied to a freshly glow-discharged grid, blotted for 3 sec with a blot force of 20 before being plunge-frozen in liquid ethane. Cryo-EM data was collected on a TITAN Krios TEM (ThermoFisher Scientific) operated at 300 KeV equipped with a 20 eV energy slit at the National Cancer Institute/NIH IRP Cryo-EM facility, Bethesda MD. Movies were recorded on a Gatan Bioquantum K3 camera as 50-frame stacks in super-resolution mode (0.415 Å per pixel) with a nominal defocus range of -0.7 to -2 µm and a total exposure dose of 54.5 e^-^ Å^-2^ over a 2.5-second exposure time.

### Cryo-EM data processing

Data processing was performed in CryoSPARC^45^. Movies were imported and corrected with a gain reference. Frame alignment of each movie was performed through Patch motion correction (multi) with the F-crop factor set at 1/2 for a pixel size of 0.83 Å per pixel. The contrast transfer function (CTF) was estimated with Patch CTF estimation (multi). Blob picker was used for initial particle picking and subsequent 2D classification. Selected good 2D classes and the corresponding particle stacks were used to train Topaz picker. All the images were re-picked with trained Topaz picker^46^, which resulted in a much larger particle stack compared to blob picker.

To eliminate bad particles, several rounds of 2D classification were performed and the particles from the good 2D classes were selected to reconstruct *ab initio* maps. Conformational flexibility was already revealed by the different NBD separations observed among the non-junk *ab initio* maps. To further sort out bad particles and conformational flexibility, multiple rounds of heterogeneous refinement and non-uniform refinement (NUR) were performed. 3D Variability Analysis (3DVA) was performed on a relatively clean particle stack after 2-3 rounds of heterogenous refinement to visualize the conformational space of DrAbcb4. For each dataset, three major conformations with distinct NBD separation distances were obtained at the end of heterogeneous and non-uniform refinements, when further refinement rounds no longer improved the resolution.

To further improve map quality, per-particle, reference-based motion correction was performed on the particle stacks of each conformation using either Bayesian polishing in RELION or Referenced-Based Motion Correction (BETA) in CryoSPARC. UCSF PyEM (https://zenodo.org/records/3576630) was used to export particle metadata from CryoSPARC to RELION. Subsequently, global and local CTF refinements were performed in CryoSPARC, followed by a final round of non-uniform refinement. Local resolution of all maps was estimated in CryoSPARC. The complete data processing workflows for the four datasets are shown in Supplementary Figures 2, 5, 6 and 7.

### Model building and refinement

ModelAngelo^47^ was used to generate the starting models automatically, which were then subjected to manual examination and adjustments in COOT^48^. The topology files for elacridar, tariquidar, and vincristine molecules were generated using PRODRG^49^. Using the two half maps, we carried out refinement in the Refmac/Servalcat package^33^ to better visualize ligand density with weighted *Fo-Fc* maps. The final models were subjected to refinement and minimization in real space using the PHENIX real-space refinement module^50^. Structure validations were performed with Molprobity^51^. The statistics of model refinement are shown in Supplementary Table 1. Structural figures were prepared using ChimeraX^52^ and PyMOL (The PyMOL Molecular Graphics System, Schrödinger).

## Data availability

The Cryo-EM density maps and atomic models generated in this study have been deposited in the Electron Microscopy Data (EMD) Bank and in the Protein Data Bank (PDB) under the following accession codes:

EMDB 70380 [https://www.ebi.ac.uk/emdb/EMD-70380] and PDB 9ODY [https://doi.org/10.2210/pdb9ODY/pdb] for Apo IF^Nar^;

EMDB 70382 [https://www.ebi.ac.uk/emdb/EMD-70382] and PDB 9OE0 [https://doi.org/10.2210/pdb9OE0/pdb] for Apo IF^Med^;

EMDB 70381 [https://www.ebi.ac.uk/emdb/EMD-70381] and PDB 9ODZ [https://doi.org/10.2210/pdb9ODZ/pdb] for Apo IF^Wide^;

EMDB 70383 [https://www.ebi.ac.uk/emdb/EMD-70383] and PDB 9OE1 [https://doi.org/10.2210/pdb9OE1/pdb] for TQR-IF^Nar^;

EMDB 70384 [https://www.ebi.ac.uk/emdb/EMD-70384] and PDB 9OE2 [https://doi.org/10.2210/pdb9OE2/pdb] for TQR-IF^Med^;

EMDB 70385 [https://www.ebi.ac.uk/emdb/EMD-70385] and PDB 9OE3 [https://doi.org/10.2210/pdb9OE3/pdb] for TQR-IF^Wide^;

EMDB 70386 [https://www.ebi.ac.uk/emdb/EMD-70386] and PDB 9OE4 [https://doi.org/10.2210/pdb9OE4/pdb] for ELA-IF^Nar^;

EMDB 70387 [https://www.ebi.ac.uk/emdb/EMD-70387] and PDB 9ODE5 [https://doi.org/10.2210/pdb9OE5/pdb] for ELA-IF^Med^;

EMDB 70388 [https://www.ebi.ac.uk/emdb/EMD-70388] and PDB 9OE6 [https://doi.org/10.2210/pdb9OE6/pdb] for ELA-IF^Wide^;

EMDB 70389 [https://www.ebi.ac.uk/emdb/EMD-70389] and PDB 9OE7 [https://doi.org/10.2210/pdb9OE7/pdb] for VCR-IF^Nar^;

EMDB 70390 [https://www.ebi.ac.uk/emdb/EMD-70390] and PDB 9OE8 [https://doi.org/10.2210/pdb9OE8/pdb] for VCR-IF^Med^;

EMDB 70391 [https://www.ebi.ac.uk/emdb/EMD-70391] and PDB 9OE9 [https://doi.org/10.2210/pdb9OE9/pdb] for VCR-IF^Wide^.

## Supporting information

suppInfo

## Acknowledgements

This research was supported by the Intramural Research Program of the National Institutes of Health (NIH), Center for Cancer Research (CCR), National Cancer Institute (NCI). All DNA sequencing services was conducted at the CCR Genomics Core, NCI. The Cryo-EM datasets were collected at the National Cancer Institute/NIH IRP Cryo-EM facility, Bethesda MD, USA. Computation for the EM image reconstruction was carried out using the Biowulf Linux cluster (biowulf.nih.gov) at the NIH high performance computing center, Bethesda, MD 20892. We thank Dr. Alexander Wlodawer for critical reading of the manuscript.

## Author Contributions Statement

J.Z. purified and characterized the protein, performed EM work and analysis, and wrote the paper; C.M.H. performed experiments, wrote the paper; L.E. analyzed data, wrote the paper; Z.C.L. & A.J.M. collected EM data. R.W.R. generated the DrAbcb4 construct; F.Z. cloned DrAbcb4 into a *Pichia* expression vector; S.V.A. wrote the paper; R.K.H. collected EM data, analyzed data, and wrote the paper. M.M.G. conceived the project and wrote the paper; D.X. conceived the project, secured funding, performed experiments, and wrote the paper.

## List of abbreviations

ABC: ATP-Binding Cassette
D_COM_: Distance between center of mass of the two NBDs
IF: inward facing
OF: outward facing
P-gp: P-glycoprotein or Permeability glycoprotein
hP-gp: human P-gp
mP-gp: murine P-gp
TM: transmembrane
TMH: TM helix

