## Supplementary material for "Functional Implications of the Conformational Landscape of a Multidrug Transporter Revealed by Structures of Zebrafish Abcb4": suppInfo

**Supplemental Information**

Supplementary Table 1

Supplementary Figures 1 to 11

**Other Supplementary Material for this manuscript includes the following:**

Supplementary Movies 1 to 2

**Table S1 | Cryo-EM data collection, model refinement and validation statistics**

| | Abcb4/Apo | | | Abcb4/Tariquidar | | | Abcb4/Elacridar | | | Abcb4/Vincristine | | |
|---|---|---|---|---|---|---|---|---|---|---|---|---|
| **Data collection**[a] | | | | | | | | | | | | |
| Total no. of micrographs | | 3,764 | | | 12,281 | | | 12,136 | | | 3,427 | |
| **Data processing** | | | | | | | | | | | | |
| Reconstruction software | | CryoSparc | | | CryoSparc/Relion | | | CryoSparc/Relion | | | CryoSparc | |
| No. extracted particles | | 1,286,409 | | | 6,064,148 | | | 5,253,173 | | | 1,841,052 | |
| No. selected 2D particles | | 880,810 | | | 2,359,321 | | | 2,498,124 | | | 540,770 | |
| Conformation designation[b] | $IF^{Nar}$ | $IF^{Med}$ | $IF^{Wide}$ | TQR-$IF^{Nar}$ | TQR-$IF^{Med}$ | TQR-$IF^{Wide}$ | ELA-$IF^{Nar}$ | ELA-$IF^{Med}$ | ELA-$IF^{Wide}$ | VCR-$IF^{Nar}$ | VCR-$IF^{Med}$ | VCR-$IF^{Wide}$ |
| No. particles for final map | 180,825 | 190,766 | 138,295 | 581,887 | 445,705 | 321,047 | 478,434 | 463,553 | 431,828 | 110,004 | 129,591 | 101,031 |
| Map resolution (Å)[c] | 3.29 | 3.39 | 3.59 | 3.02 | 3.18 | 3.54 | 3.00 | 3.16 | 3.31 | 3.95 | 3.70 | 4.09 |
| Map sharpening B-factor ($Å^2$) | -116.0 | -116.9 | -114.6 | -126.2 | -110.1 | -129.9 | -109.9 | -116.9 | -110.9 | -155.5 | -129.2 | -178.1 |
| **PHENIX refinement statistics** | | | | | | | | | | | | |
| Initial model used | PDB:5KO2 | PDB:5KO2 | PDB:5KO2 | ModelAngelo | ModelAngelo | ModelAngelo | ModelAngelo | ModelAngelo | ModelAngelo | ModelAngelo | ModelAngelo | ModelAngelo |
| Model composition | | | | | | | | | | | | |
| Non-hydrogen Atoms | 9,010 | 9,010 | 8,978 | 9,322 | 9,267 | 9,001 | 9,273 | 9,044 | 8,984 | 9,243 | 9,034 | 9,013 |
| No. of protein residues | 1,154 | 1,154 | 1,150 | 1,178 | 1,176 | 1,152 | 1,179 | 1,159 | 1,151 | 1,176 | 1,157 | 1,155 |
| No. of Ligands | $Mg^{2+}$: 2<br>ATPγS: 2 | $Mg^{2+}$: 2<br>ATPγS: 2 | $Mg^{2+}$: 2<br>ATPγS: 2 | $Mg^{2+}$: 2<br>ATPγS: 2<br>TQR: 2 | $Mg^{2+}$: 2<br>ATPγS: 2<br>TQR: 1<br>CLR[e]: 1 | $Mg^{2+}$: 2<br>ATPγS: 2 | $Mg^{2+}$: 2<br>ATPγS: 2<br>ELA: 1<br>CHD[e]:1 | $Mg^{2+}$: 2<br>ATPγS: 2 | $Mg^{2+}$: 2<br>ATPγS: 2 | $Mg^{2+}$: 2<br>ATPγS: 2<br>VCR: 1 | $Mg^{2+}$: 2<br>ATPγS: 2 | $Mg^{2+}$: 2<br>ATPγS: 2 |
| Overall B-factor ($Å^2$) | | | | | | | | | | | | |
| Protein | 67.79 | 68.48 | 97.68 | 54.30 | 61.88 | 73.48 | 112.4 | 107.5 | 137.4 | 205.8 | 167.5 | 260.9 |
| Ligands | 102.81 | 98.52 | 124.02 | 70.43 | 85.66 | 95.13 | 133.6 | 135.9 | 160.7 | 260.1 | 196.6 | 290.6 |
| RMS deviations | | | | | | | | | | | | |
| Bond length (Å) | 0.005 | 0.003 | 0.005 | 0.003 | 0.004 | 0.005 | 0.003 | 0.003 | 0.003 | 0.007 | 0.003 | 0.004 |
| Bond angle (º) | 0.663 | 0.554 | 1.068 | 0.677 | 0.965 | 0.977 | 0.681 | 0.657 | 0.598 | 1.229 | 0.672 | 0.964 |
| Model resolution[d] (Å) | 3.72 | 3.77 | 3.99 | 3.28 | 3.36 | 3.84 | 3.28 | 3.38 | 3.54 | 4.31 | 4.03 | 4.54 |
| Real-space correlation | | | | | | | | | | | | |
| CC (mask) | 0.80 | 0.79 | 0.79 | 0.83 | 0.85 | 0.74 | 0.88 | 0.85 | 0.82 | 0.87 | 0.83 | 0.85 |
| CC (volume) | 0.79 | 0.78 | 0.78 | 0.82 | 0.84 | 0.73 | 0.87 | 0.84 | 0.81 | 0.87 | 0.83 | 0.85 |
| CC (peaks) | 0.66 | 0.66 | 0.58 | 0.72 | 0.72 | 0.58 | 0.64 | 0.60 | 0.52 | 0.58 | 0.52 | 0.53 |
| **Validation** | | | | | | | | | | | | |
| MolProbity score | 2.05 | 2.33 | 2.40 | 2.20 | 2.36 | 1.99 | 2.02 | 1.57 | 1.96 | 2.64 | 1.69 | 1.80 |
| Clash score | 5.89 | 8.09 | 10.39 | 9.57 | 8.85 | 4.53 | 4.81 | 3.35 | 8.95 | 20.15 | 4.89 | 7.98 |
| Rotamer outliers (%) | 2.62 | 3.04 | 2.73 | 2.86 | 4.10 | 3.05 | 3.98 | 1.67 | 1.58 | 4.00 | 1.25 | 0.00 |
| Ramachandran plot | | | | | | | | | | | | |
| Favored (%) | 93.82 | 90.94 | 90.56 | 95.05 | 93.68 | 94.07 | 95.48 | 96.01 | 95.28 | 94.44 | 94.70 | 94.69 |
| Allowed (%) | 5.92 | 8.71 | 9.00 | 4.78 | 6.15 | 5.76 | 4.43 | 3.90 | 4.72 | 5.56 | 5.21 | 5.22 |
| Disallowed (%) | 0.26 | 0.35 | 0.44 | 0.17 | 0.17 | 0.17 | 0.09 | 0.09 | 0.00 | 0.00 | 0.09 | 0.09 |
| **Data deposition** | | | | | | | | | | | | |
| PDB | 9ODY | 9OE0 | 9ODZ | 9OE1 | 9OE2 | 9OE3 | 9OE4 | 9OE5 | 9OE6 | 9OE7 | 9OE8 | 9OE9 |
| EMD | EMD-70380 | EMD-70382 | EMD-70381 | EMD-70383 | EMD-70384 | EMD-70385 | EMD-70386 | EMD-70387 | EMD-70388 | EMD-70389 | EMD-70390 | EMD-70391 |

a - All data sets were collected on a Thermo Fisher Scientific Titan Krios microscope (accelerating voltage of 300kV) equipped with a Gatan K3 camera at a nominal magnification of 105,000x and a pixel size of 0.415 Å (0.83Å binned). Each movie was dose fractionated into 50 frames with a total electron exposure of 54.4 $e^-/Å^2$ at a defocus range of 0.7-2.0 μm.

b - Conformation designation: Nar - narrow separation between two NBDs; Med - medium separation; Wide - wide separation.

c - map resolution was based on FSC threshold of 0.143.

d - model resolution is defined as the FSC of 0.5 for model vs. map

e - CLR, cholesterol; CHD, cholate.

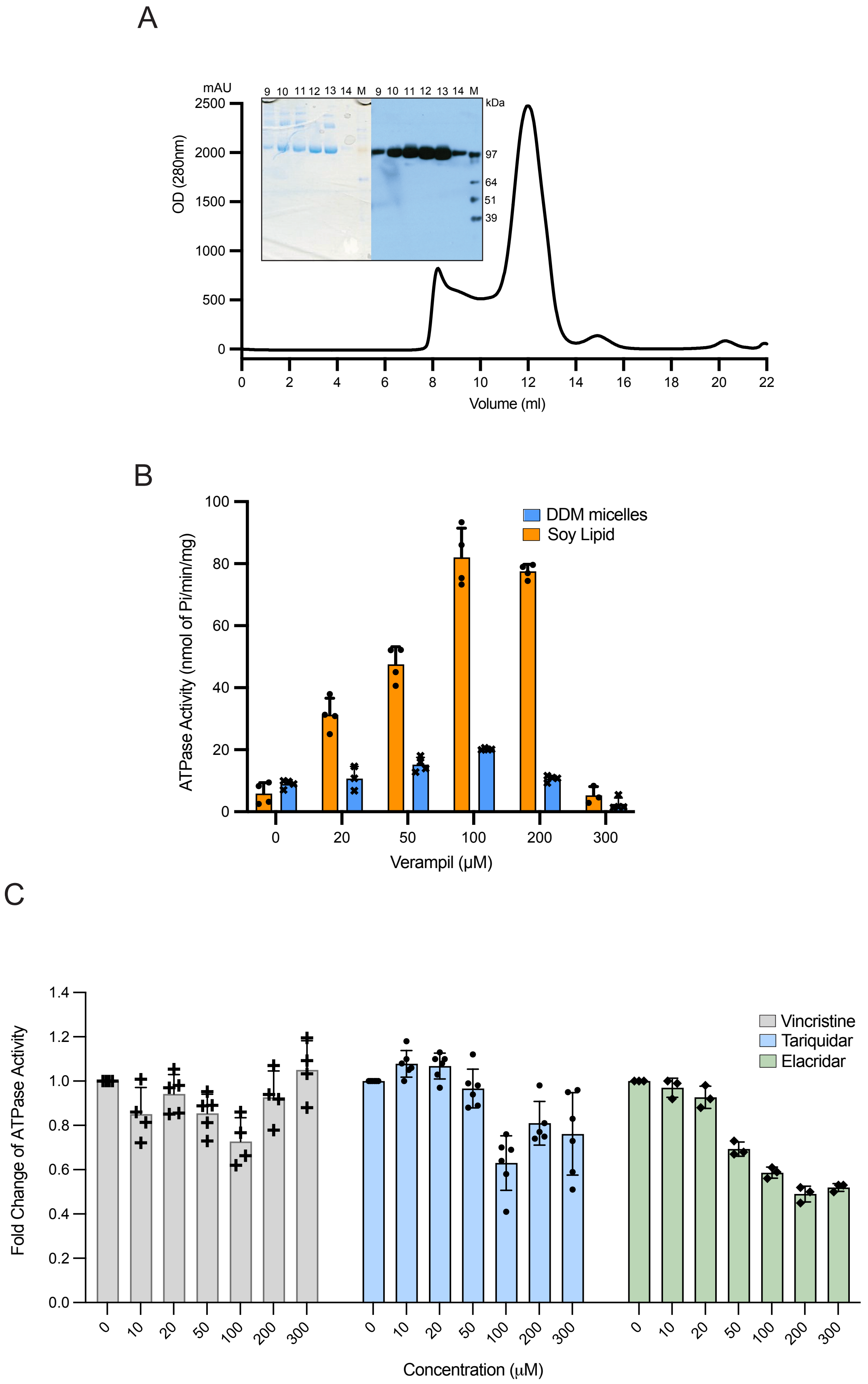

Zhan et al., Figure S1

**Figure S1. Expression, purification, biochemical characterization of recombinant DrAbcb4.** (A) Size-exclusion chromatogram (SEC) of isolated DrAbcb4 using a Superdex$^{TM}$ 200 Increase 10/300 GL column. Inset: SDS-PAGE (left) and Western Blot (right) analyses of SEC peak fractions showing monodispersity of purified sample and cross-reaction of the monoclonal antibody C219 with DrAbcb4. (B) Basal and verapamil-modulated ATPase activity in detergent and in soy lipid bicelle solutions. (C) Concentration-dependent ATPase activity of DrAbcb4 in the presence of vincristine, tariquidar or elacridar.

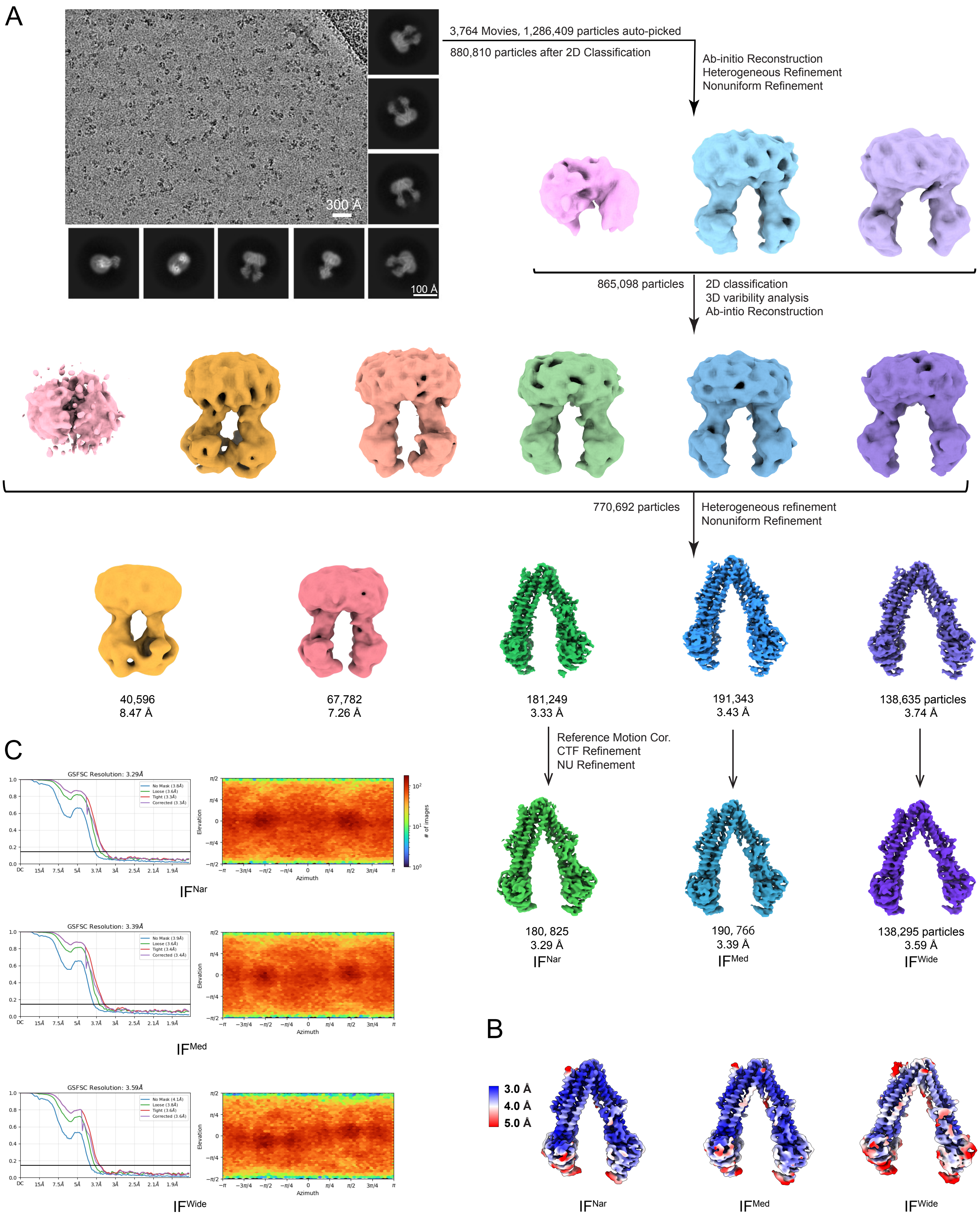

Zhan et al., Figure S2

**Figure S2. Cryo-EM data processing for DrAbcb4 in the presence of ATPγS/$Mg^{2+}$.** (A) Processing workflow included particle picking, 2D classification, 3D reconstruction and classification that ultimately led to three high-resolution structures of DrAbcb4 in IF conformations. (B) Local resolution maps for the three final structures. (C) Gold-standard Fourier shell correlation (FSC) curves (left) and viewing direction distribution plots (right) for the three reconstructions.

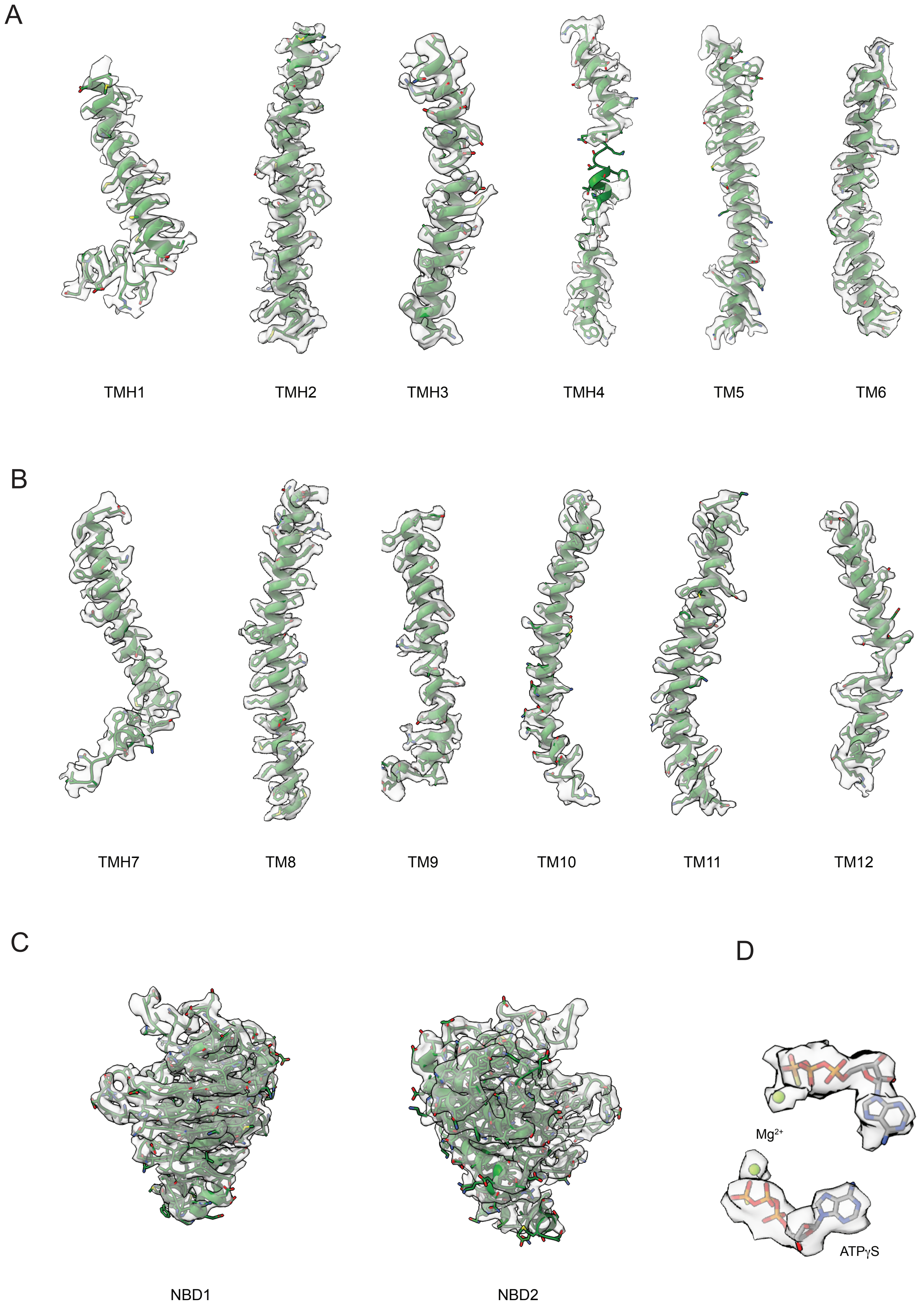

Zhan et al., Figure S3

**Figure S3. Representative EM density for various regions of DrAbcb4.** Selected regions of the sharpened EM density map of DrAbcb4 in the Apo $IF^{Nar}$ conformation are shown as semi-transparent surfaces contoured at a level of 0.018, overlaid with the corresponding atomic models. Displayed regions are: (A) TMHs 1-6, (B) TMHs 7-12, (C) NBD1 and NBD2, and (D) bound ATP$\gamma$S.

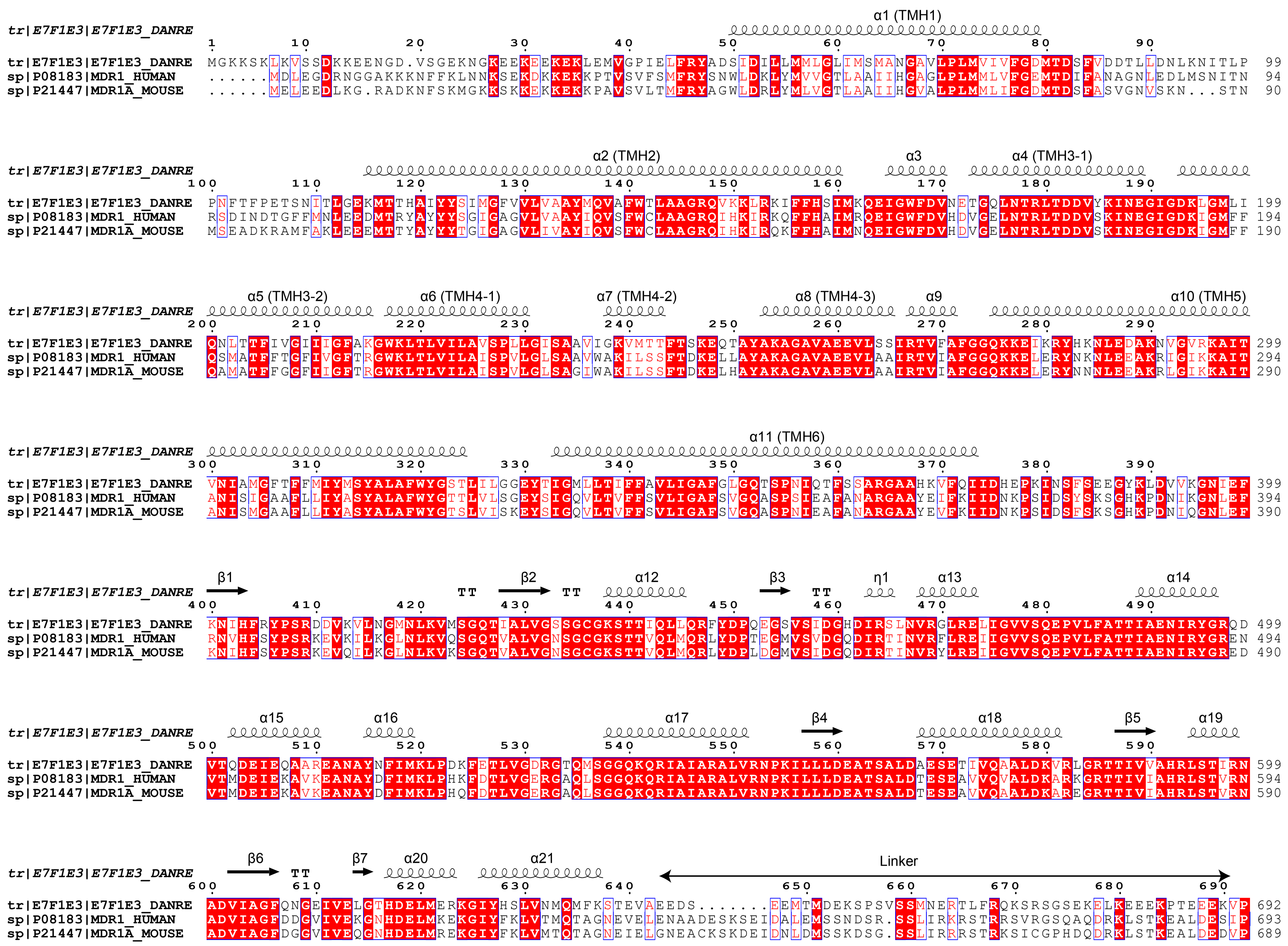

Zhan et al., Figure S4A

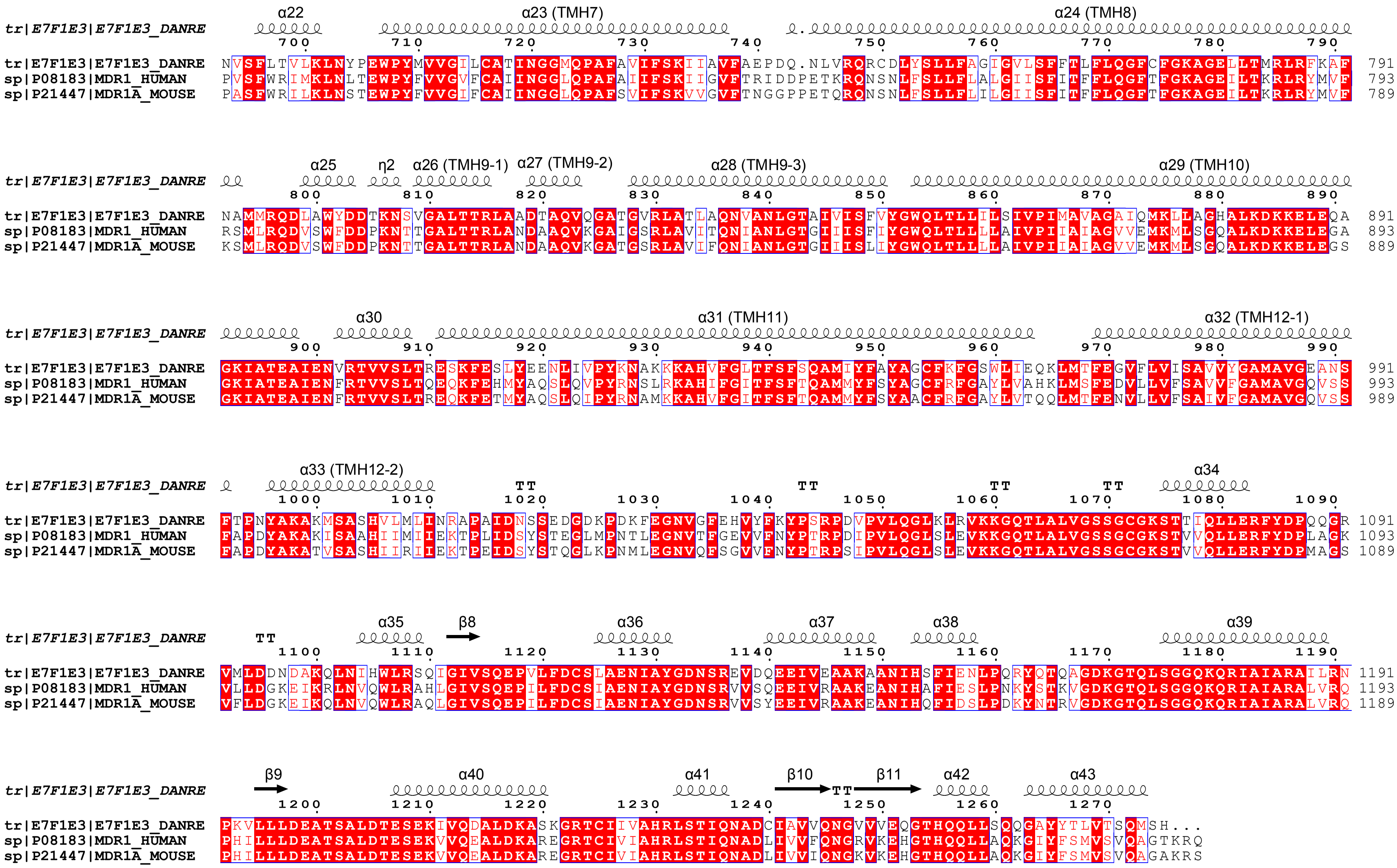

Zhan et al., Figure S4B

**Figure S4. Sequence alignment of zebrafish Abcb4, hP-gp, and mP-gp.** Secondary structure assignments based on Apo Abcb4 structure are placed above the alignment (A) N-terminal half. (B) C-terminal half.

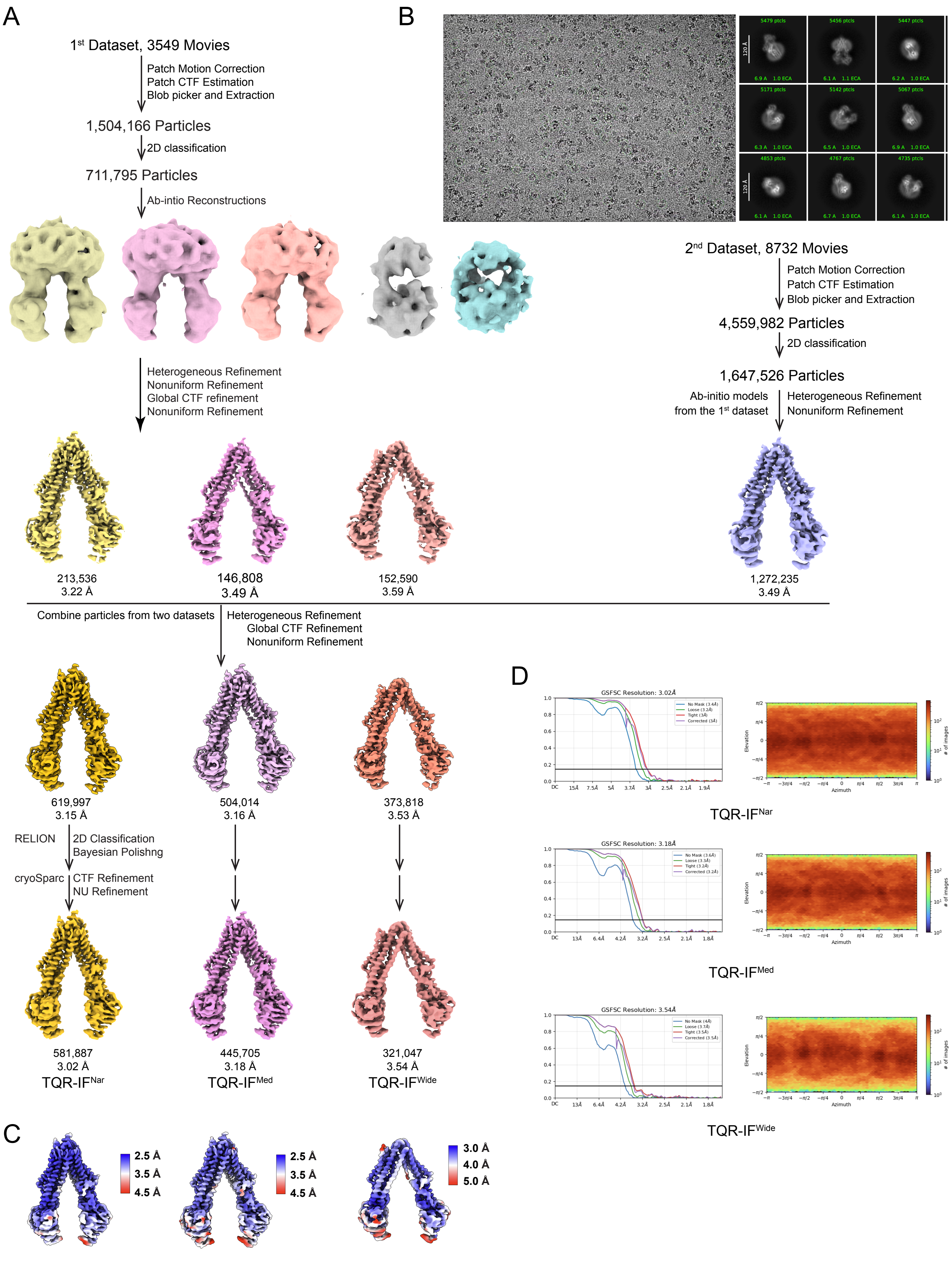

Zhan et al., Figure S5

**Figure S5. Cryo-EM data processing workflow for DrAbcb4/tariquidar.** The final dataset includes (A) subset 1 with 3,549 movies and (B) subset 2 with 8,732 movies, collected from the same DrAbcb4 sample incubated with tariquidar, ATPγS, and $MgCl_2$. Representative raw micrograph and 2D class averages are given for the 2nd data set. Particles from the two data sets were combined. 3D classifications and reconstructions obtained are shown. (C) Final 3D maps of the three major conformations, colored by local resolution. (D) Gold-standard FSC curves (left) and viewing direction distribution plots (right) for the three final reconstructions.

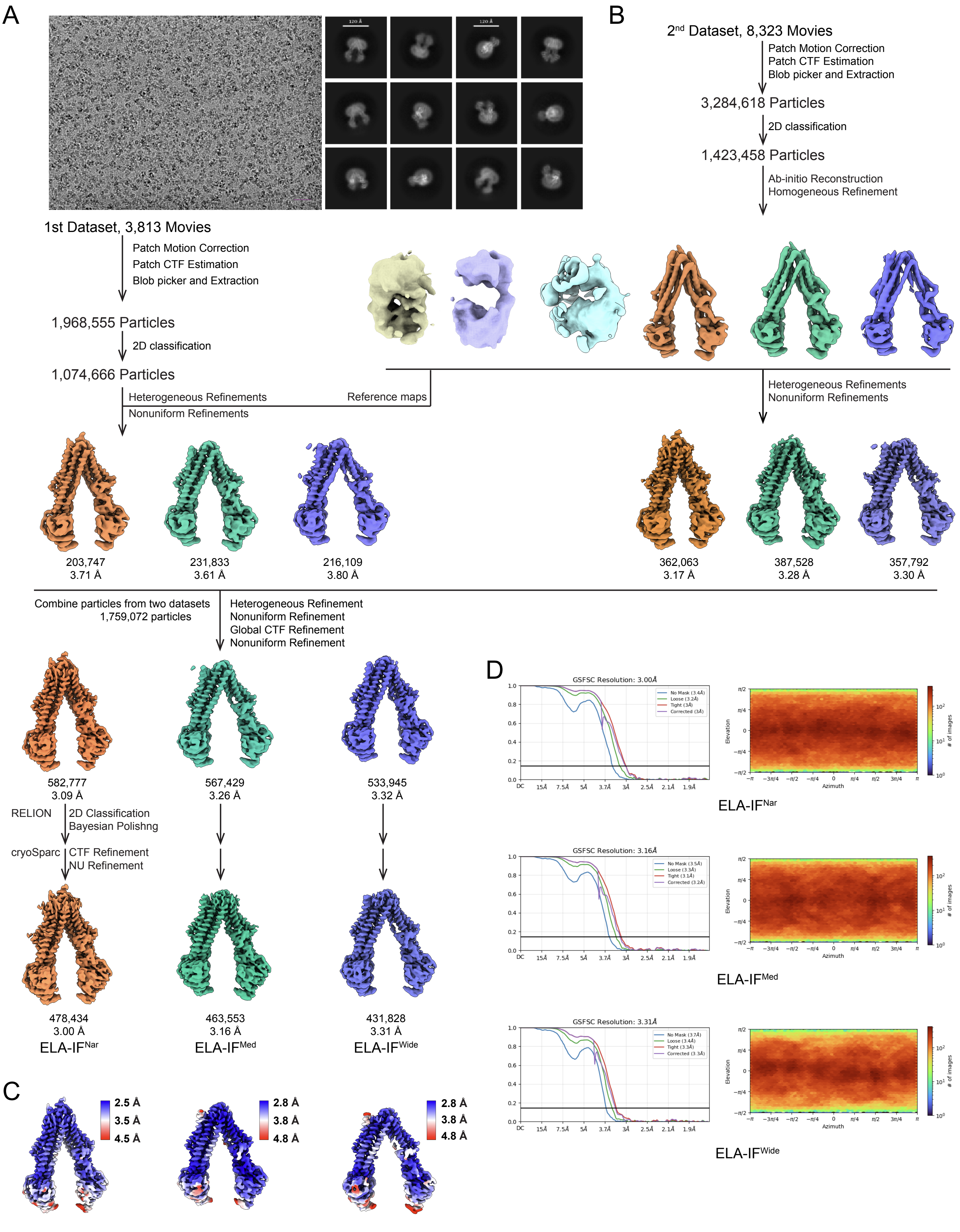

Zhan et al., Figure S6

**Figure S6. Cryo-EM data processing workflow for DrAbcb4/elacridar dataset.** The final dataset comprises (A) a smaller subset with 3,813 movies and (B) a larger subset with 8,323 movies, collected from the same DrAbcb4 sample incubated with elacridar, ATP$\gamma$S, and $MgCl_2$. Representative micrograph and 2D class averages for the 1st data set are shown. Particles from the two data sets were combined. 3D reconstructions from the data processing workflow are shown. (C) Final 3D maps of the three major conformations, colored by local resolution. (D) Gold-standard FSC curves (left) and viewing direction distribution plots (right) for the three final reconstructions.

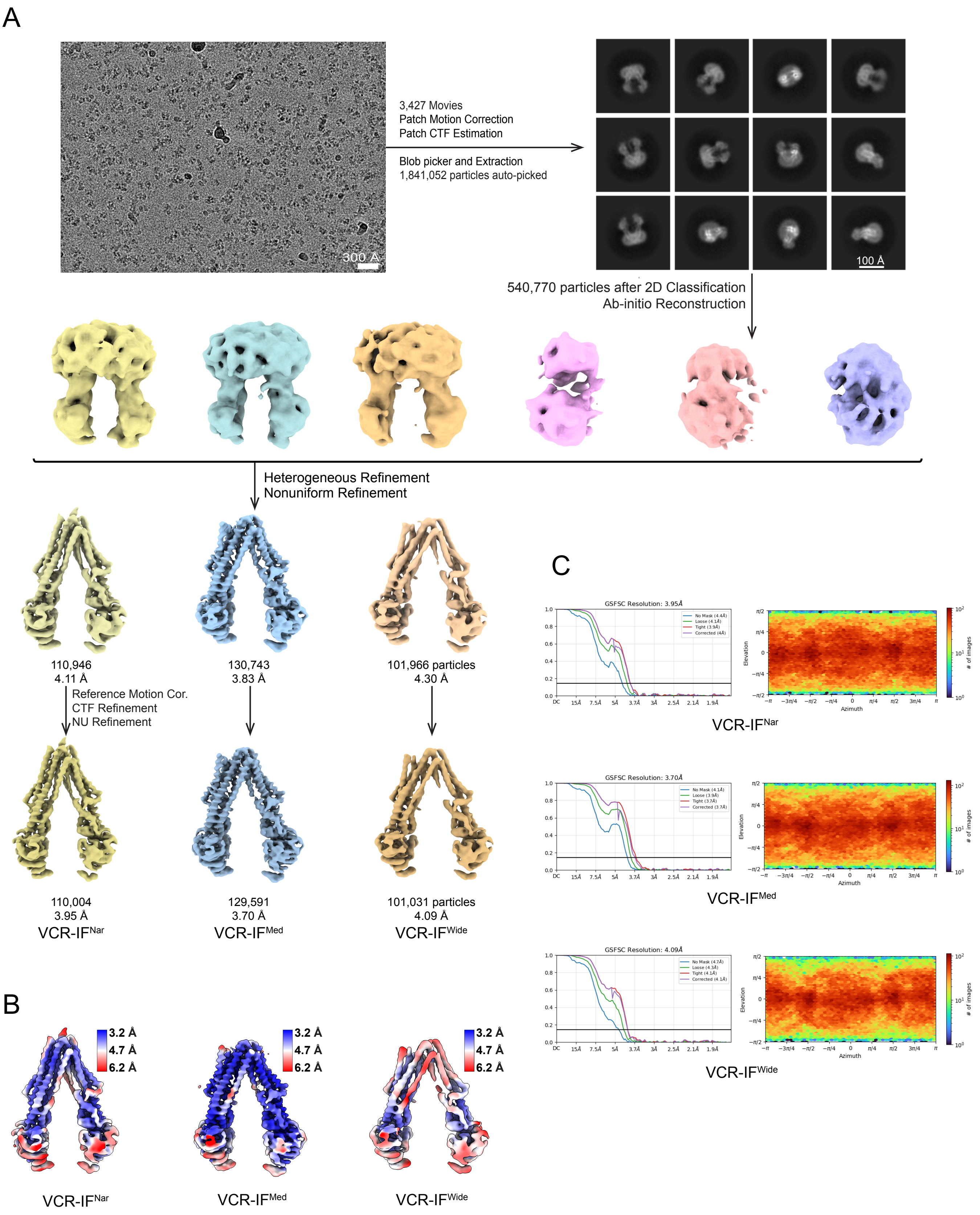

Zhan et al., Figure S7

**Figure S7. Cryo-EM data processing workflow for DrAbcb4/vincristine dataset.** (A) Representative micrograph, 2D class averages, and 3D reconstructions from the DrAbcb4/vincristine dataset are shown. Three major conformations with resolutions between 3.7 Å to 4.1 Å were obtained. (B) Final 3D reconstructions of the three conformations, colored by local resolution. (C) Gold-standard FSC curves (left) and viewing direction distribution plots (right) for the three final reconstructions.

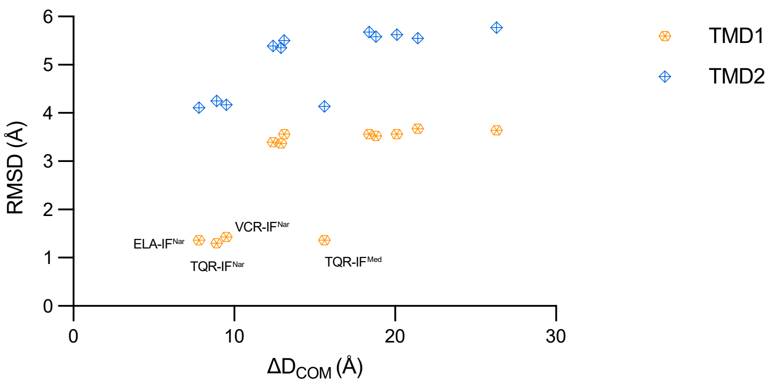

**Figure S8. Structural deviations of mP-gp and DrAbcb4 as a function of $D_{COM}$.** Using mP-gp (PDB:5KPD) as a reference for superposing the 12 DrAbcb4 structures pairwise, the rms deviations (RMSD) are plotted against the changes in $D_{COM}$ ($\Delta D_{COM}$), revealing a positive correlation indicative of coordinated movement.

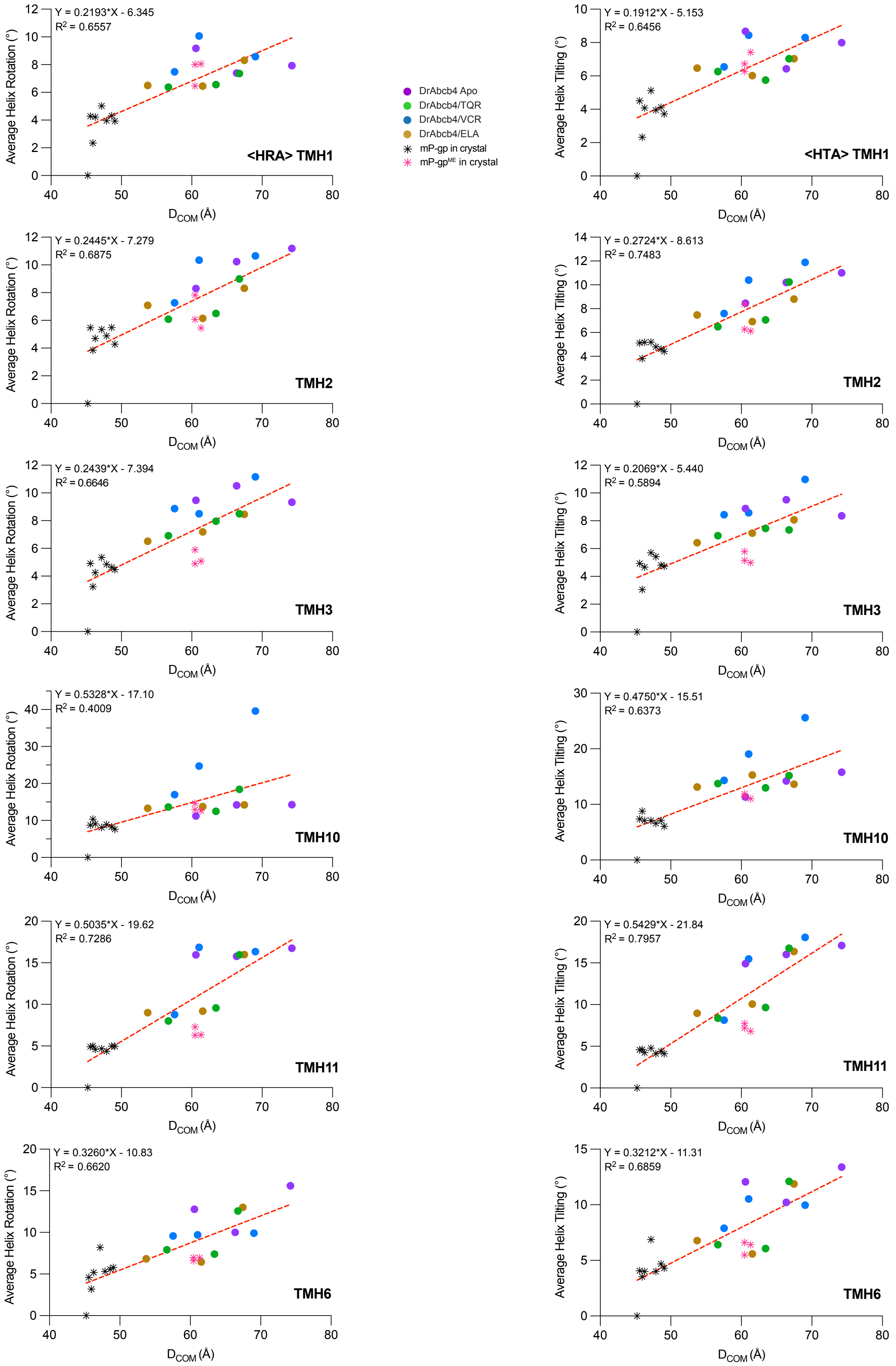

Zhan et al., Figure S9A

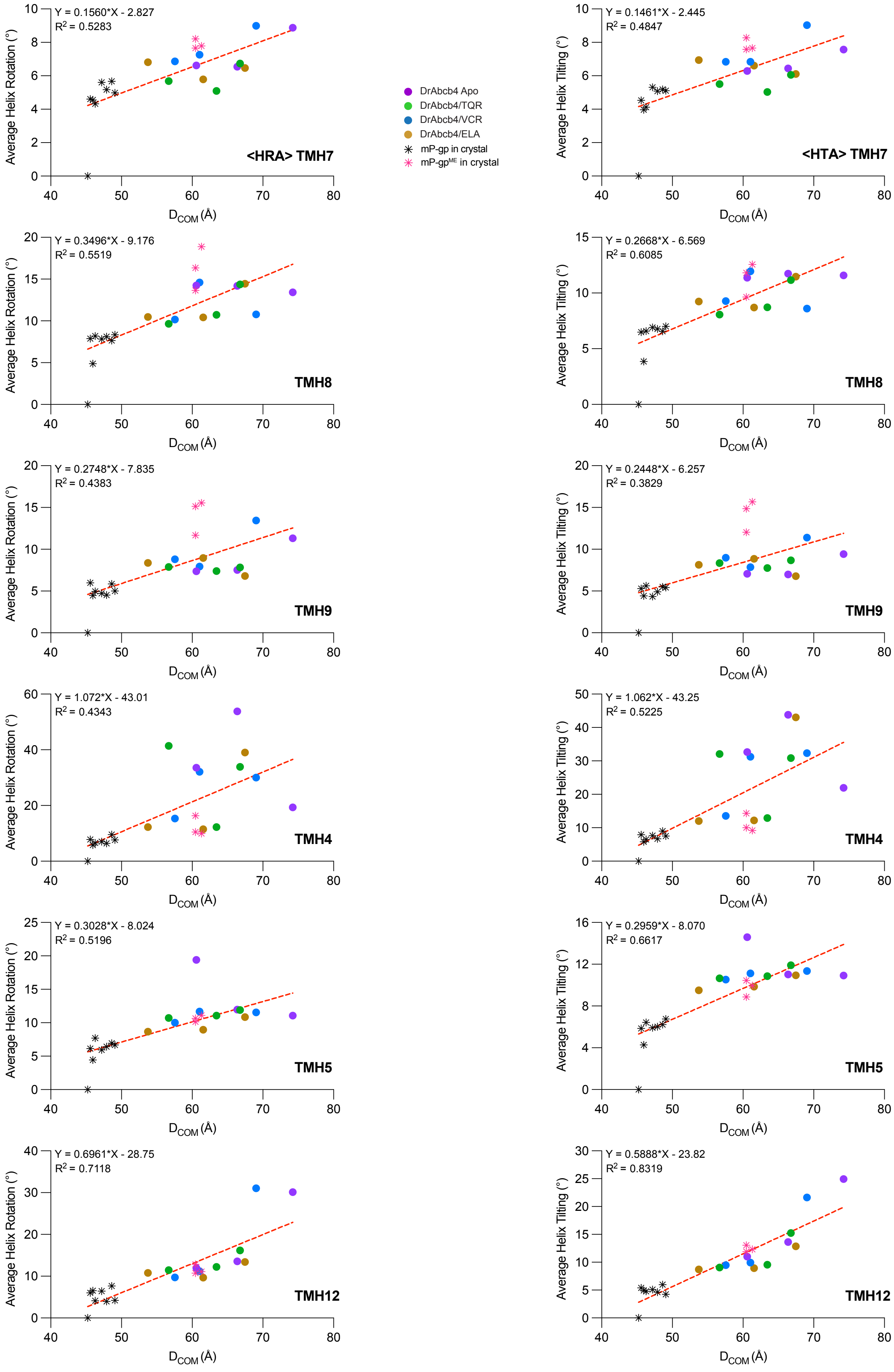

Zhan et al., Figure S9B

**Figure S9. Scatter plots showing the correlation of $D_{COM}$ with average rotation angle <HRA> and average tilting angle <HTA> of individual TM helices in TMD1 (Figure S9A) and in TMD2 (Figure S9B).** Left: Scatter plot of <HRA> as a function of $D_{COM}$. Left: Scatter plot of <HTA> as a function of $D_{COM}$. The dashed lines are trending lines through the sample dots. Fitting equations and R values are also given.

| TMD1 | hP-gp | mP-gp | DrAbcb4 |
|---|---|---|---|
| TM1 | L65 | L64 | L70 |
| TM1 | M68 | M67 | M73 |
| TM1 | M69 | M68 | V74 |
| TM1 | F72 | F71 | F77 |
| TM6 | F336 | F332 | F341 |
| TM6 | V338 | V334 | V343 |
| TM6 | L339 | L335 | L344 |
| TM6 | I340 | I336 | I345 |
| TM6 | A342 | A338 | A347 |
| TM6 | F343 | F339 | F348 |
| TM6 | S344 | S340 | G349 |
| * TM6 | G346 | G342 | G351 |
| TM6 | Q347 | Q343 | Q352 |
| TM6 | P350 | P346 | P355 |
| TM10 | E875 | E871 | Q873 |
| TM10 | M876 | M872 | M874 |
| TM10 | L879 | L875 | L877 |
| TM10 | S880 | S876 | A878 |
| * TM11 | T945 | T941 | S943 |
| TM11 | Q946 | Q942 | Q944 |
| TM11 | M949 | M945 | I947 |
| TM11 | Y950 | Y946 | Y948 |
| TM11 | Y953 | Y949 | Y951 |

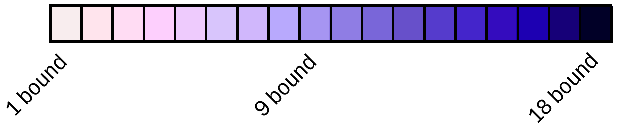

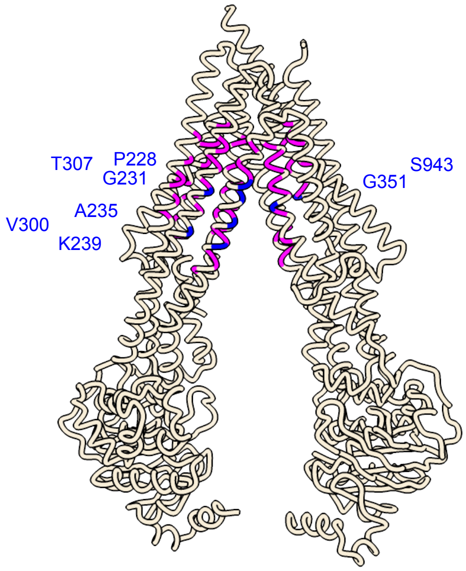

| | DrAbcb4 | mP-gp | hP-gp | TMD2 | |
|---|---|---|---|---|---|
| | N720 | N717 | N721 | TM7 | |
| | Q724 | Q721 | Q725 | TM7 | |
| | F727 | F724 | F728 | TM7 | |
| | A728 | S725 | A729 | TM7 | |
| | F731 | F728 | F732 | TM7 | |
| | F757 | F755 | F759 | TM8 | |
| | G761 | G759 | G763 | TM8 | |
| | S764 | S762 | S766 | TM8 | |
| | F765 | F763 | F767 | TM8 | |
| | L768 | F766 | F770 | TM8 | |
| | Q771 | Q769 | Q773 | TM8 | |
| | V829 | S827 | S831 | TM9 | |
| | A832 | A830 | A834 | TM9 | |
| | Q836 | Q834 | Q838 | TM9 | |
| | I223 | I214 | I218 | TM4 | |
| | V226 | I217 | I221 | TM4 | |
| | S227 | S218 | S222 | TM4 | |
| | P228 | P219 | P223 | TM4 | * |
| | L230 | L221 | L225 | TM4 | |
| | G231 | G222 | G226 | TM4 | * |
| | A234 | A225 | A229 | TM4 | |
| | A235 | G226 | A230 | TM4 | * |
| | I237 | W228 | W232 | TM4 | |
| | G238 | A229 | A233 | TM4 | |
| | K239 | K230 | K234 | TM4 | * |
| | M241 | L232 | L236 | TM4 | |
| | T242 | S233 | S237 | TM4 | |
| | F244 | F235 | F239 | TM4 | |
| | V300 | A291 | A295 | TM5 | * |
| | M304 | M295 | I299 | TM5 | |
| | T307 | A298 | A302 | TM5 | * |
| | F308 | F299 | F303 | TM5 | |
| | I311 | I302 | I306 | TM5 | |
| | Y312 | Y303 | Y307 | TM5 | |
| | S314 | S305 | S309 | TM5 | |
| | Y315 | Y306 | Y310 | TM5 | |
| | I976 | F974 | F978 | TM12 | |
| | S977 | S975 | S979 | TM12 | |
| | Y981 | F979 | F983 | TM12 | |
| | M984 | M982 | M986 | TM12 | |
| | A985 | A983 | A987 | TM12 | |
| | E988 | Q986 | Q990 | TM12 | |
| | A989 | V987 | V991 | TM12 | |
| | F992 | F990 | F994 | TM12 | |

Zhan et al., Figure S10

**Figure S10. Sequence conservation for residues interacting with bound compounds.** The left panel lists all drug-interacting residues from TM helices of TMD1, and the right panel shows those from TM helices of TMD2. Corresponding residues in hP-gp, mP-gp, and DrAbcb4 sequences are given with colored squares indicating the frequency of each residue found interacting with bound compounds. Newly identified drug-interacting residues in DrAbcb4 structures are marked with an asterisk, and their locations are highlighted in blue in the ribbon diagram of DrAbcb4.

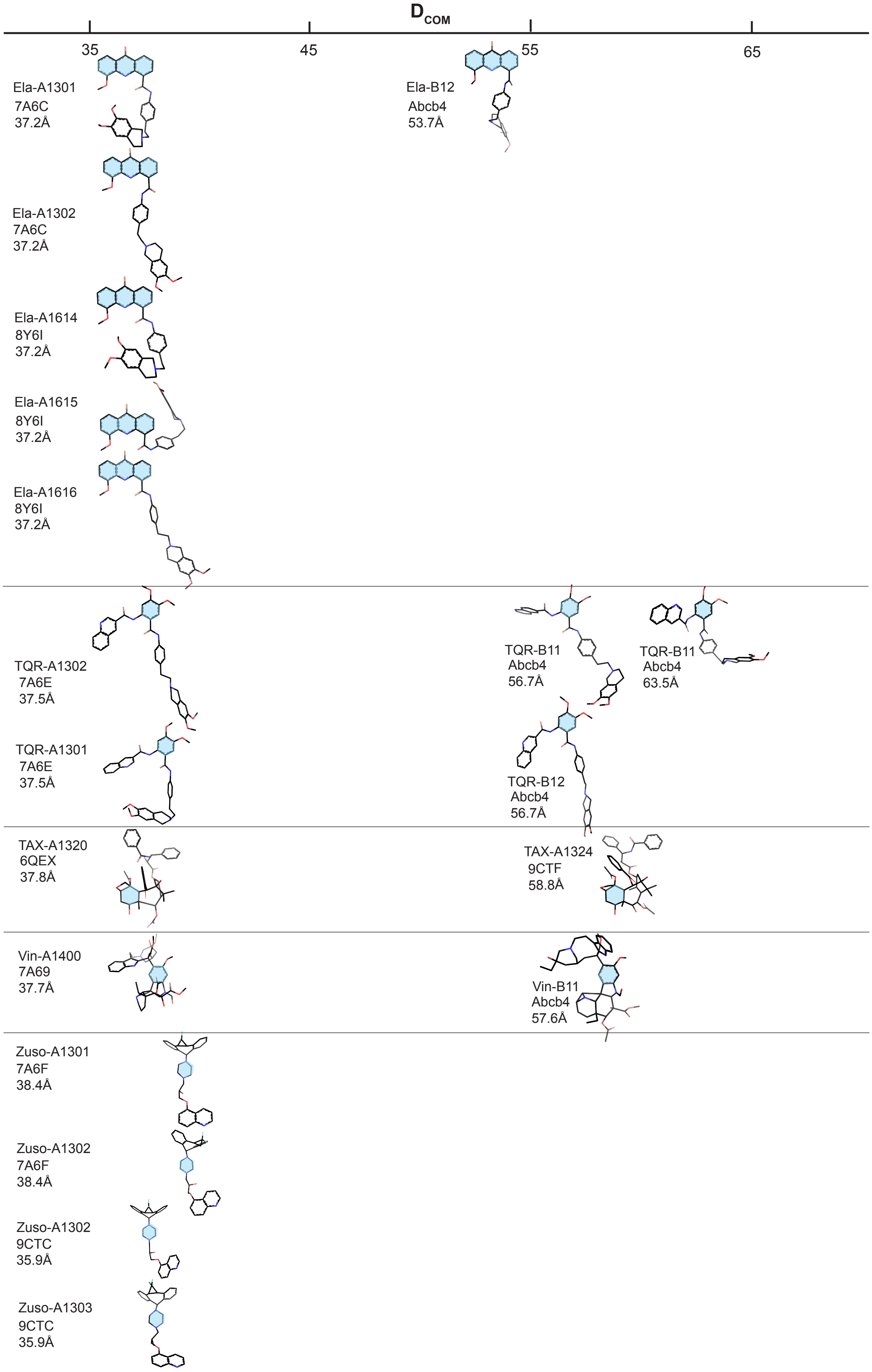

Zhan et al., Figure S11

**Figure S11. Poses of bound compounds arranged by $D_{COM}$ values.** Compounds are illustrated as stick models and labelled with their PDB codes and $D_{COM}$ values. The chemical moieties that were used for alignment are colored cyan.

**Legends for supplementary movies**

**Movie S1. 3D Variability Analysis for the DrAbcb4 apo dataset**

**Movie S2. 3D Variability Analysis for the DrAbcb4/Tariquidar dataset**
